# miR-10b-5p regulates adipocyte lineage commitment and adipogenesis via targeting of Gata6 and Tubby

**DOI:** 10.1101/2025.07.12.664409

**Authors:** Nikoletta Kalenderoglou, Federica Dimitri, Carmen Navarro Gonzalez, Antonio Vidal-Puig, Jacob Hobbs, Awais Younis, Stefania Carobbio, Mark Christian

## Abstract

2

**Background:** Adipogenesis is a highly organised series of events that facilitates the healthy expansion of adipose tissue, beginning during embryogenesis and continuing throughout life. White adipogenesis protects against lipotoxicity, influencing insulin resistance and obesity-related comorbidities. Brown adipogenesis enhances energy expenditure, thereby counteracting weight gain, lipotoxicity and insulin resistance. Recently, there has been a significant increase in interest regarding adipocyte differentiation, mainly focusing on the interplay between microRNAs (miRNAs) and the transcriptional cascade that governs adipogenesis and metabolic dysfunction. This study aimed to identify miRNAs regulating white and brown adipocyte differentiation and define miRNA action in a stem cell model of adipogenesis.

**Methods:** Small RNAseq analysis of primary mouse brown and white adipocytes (WAs) identified miR-10b to be upregulated in mature brown adipocytes (BAs). We generated two model systems: 1) immortalized brown pre-adipocytes treated with miRNA inhibitors and 2) CRISPR/Cas9 KO of miR-10b in E14 mouse embryonic stem cells (mESCs). Both cell models were differentiated into mature adipocytes. To unravel the pathways that are affected by miR-10b depletion, a transcriptomic analysis was performed at key time points.

**Results:** Both cell models showed that miR-10b-5p depletion severely impaired differentiation into mature adipocytes, as indicated by a lack of lipid droplet formation and reduced adipogenic gene expression. Gene expression analysis supports that miR-10b-5p directs embryonic stem (ES) cells towards the mesoderm lineage, promoting commitment to pre-adipocytes by downregulating Gata6 and its downstream target Bmp2. This mechanism appears to be unaffected in BAs. Our study demonstrated that miR-10b-5p regulates the later stages of adipogenesis, at least in part, by downregulating Tub, a direct target of miR-10b-5p. We also confirmed that miR-10b-5p alleviated the halted differentiation phenotypes of adipocytes by supressing the G Protein Signalling pathway mediated by Tubby.

**Conclusions:** These results evidence that miR-10b inhibition plays a dynamic role in adipocyte biology, as its inhibitory effects manifest differently during the stem cell preadipocyte proliferation state and during the maturation phase of adipocytes. Collectively, our study demonstrated that miR-10b-5p may represent a new potential therapeutic target for lipodystrophy and obesity.

## 4 Background

Adipogenesis is a coordinated process regulated by multiple autocrine, paracrine and endocrine signals. It facilitates the healthy expansion of adipose tissue, beginning during embryogenesis and continuing throughout life (1). It is generally regarded as a two-step process: a commitment step, wherein a fibroblast-like mesenchymal stem cell (MSC) commits to the adipocyte lineage without any morphological alterations, developing into a preadipocyte. This commitment is then followed by differentiation, wherein committed preadipocytes expand, undergo growth arrest, induced by contact inhibition, and take on the characteristics of functional mature adipocytes as they acquire the machinery that is essential for lipid transport and synthesis, insulin sensitivity and the secretion of adipocyte-specific proteins (2).

In mammals, two major types of adipose tissue exist, white (WAT) and brown (BAT). WAT plays a critical role in the regulation of energy homeostasis by storing excess energy in the form of triglycerides (TGs), secreting adipokines that regulate lipid metabolism, secreting inflammatory cytokines, and controlling insulin sensitivity (3,4). The lipids in adipose tissue are produced by *de novo* lipogenesis, and the hydrolysis and intracellular re-esterification of TGs from circulating lipoproteins. They serve not only as a caloric reservoir for future need but also as a protective agency in cells to sequester lipids away from peripheral tissues that are susceptible to lipotoxicity, which disrupts insulin action on metabolism (5). In contrast to WAT, BAT is composed of multiple smaller (multilocular) lipid droplets. Classical BAT is characterised as a temperature mediator organ specialised for oxidizing fatty acids and glucose to produce heat through non-shivering thermogenesis (6,7).

Adipocytes, like other mesenchymal cells, emerge from the mesodermal layer of the embryo. Specifically, white adipocytes (WAs) are believed to derive from the lateral plate mesoderm, whereas brown adipocytes (BAs) are developed from paraxial mesoderm (8). Furthermore, MSCs can be committed to either an adipogenic lineage of myogenic factor 5 (Myf5)-negative cells or a myogenic lineage of Myf5-positive cells (9). Previous studies using lineage tracing experiments in mouse models have demonstrated that white adipocytes emerge from the adipogenic lineage, whereas cells expressing Myf5 give rise to skeletal myocytes and brown adipocytes in interscapular and perirenal depots (10). Improved knowledge of adipogenesis is necessary to gain insight into brown and white fat physiology. Interest in adipocyte differentiation has increased markedly over the past few years, with emphasis on the intersection between miRNAs and the transcriptional cascade that controls adipogenesis and metabolic dysfunction.

miRNAs are endogenous, noncoding small RNAs that play a key role in regulating various developmental and cellular functions. They control gene expression at the posttranscriptional level by binding their seed sequence to complementary regions in the 3’ untranslated region (UTR) of target mRNAs, leading to either the destabilization of the mRNA or the suppression of its translation (11,12). miRNA targets are highly tissue- or cell type-dependent, and a single miRNA can perform diverse functions by controlling different mRNAs in different tissues (11,13). In search of miRNAs crucial for adipogenesis, we identified miR-10b-5p as a novel regulator through small RNA sequencing of mouse BAs and WAs. miR-10b-5p was upregulated in BAs, and its depletion in immortalized brown pre-adipocyte and stem cell models impaired differentiation into mature adipocytes. Our findings strongly suggest that miR-10b depletion impairs the early stages of adipogenesis, specifically the commitment of ESCs to preadipocytes. This effect may be driven by directing cells toward the mesoderm lineage through the upregulation of GATA6 and its downstream target, BMP2. Notably, this mechanism appears unaffected in brown adipocytes. RNA sequencing revealed that ablation of miR-10b-5p led to a significant increase in genes related to G Protein signalling associated with elevated Tubby (Tub) which was identified as direct target of miR-10b-5p. The suppression in adipocyte differentiation after miR-10b-5p inhibition was partially reversed by Tub silencing. Our research highlights the pleiotropic regulatory role of miR-10b-5p during adipogenesis.

## 5 Methods

### 5.1 Cell culture

Primary cultures of subcutaneous white adipose tissue (scWAT) and interscapular brown adipose tissue (iBAT) were collected from 7-day-old C57BL/6 mice. scWAT was digested in DMEM-OK, which consists of DMEM/F12 (Sigma) supplemented with 2% BSA, 16 µM biotin (B4501, Sigma), 1.8 µM pantothenic acid (24330100, Acros Organics), and 100 µM ascorbic acid (A8960, Sigma), along with 2 mg/ml collagenase A (Sigma). The BAT was digested using a digestion buffer composed of 100 mM HEPES (pH 7.4), 123 mM NaCl, 5 mM KCl, 3 mM CaCl₂, 5 mM glucose, 1.5% BSA, and 1 mg/ml collagenase A in DMEM-OK. The mixture was incubated with agitation at 37°C for 30 minutes. The digested tissue was then filtered and centrifuged to isolate the stromal vascular fraction (SVF). The SVF pellet from scWAT was resuspended and treated with a lysis buffer containing 100 mL of Solution A (2.08 g TRIS, pH 7.65, in 100 mL distilled H_2_O) and 900 mL of Solution B (8.3 g NH_4_Cl in 1000 mL distilled H_2_O), before centrifugation again. The scWAT cell pellet was resuspended in DMEM-OK supplemented with 10% NCS. Cells from scWAT were then plated in a 6-well plate, maintained till confluency and differentiated in DMEM-OK with 10% NCS, supplemented with 5 mg/ml insulin, 33.33 nM dexamethasone (D2915, Sigma), 1mg/ml transferrin (03-0124SA, Gibco), and 2ng/ml T3 (T6397, Sigma). Media was changed every 2–3 days, with differentiation occurring within a week. The SVF pellet from iBAT was washed once more in DMEM-OK, centrifugated for 5 min at 2300 rpm and plated in 6 well plates. Details of the induction and maintenance media for brown adipocytes are presented in Supplementary Table 1. Small RNA-Seq was performed with RNA from primary white and brown pre-adipocytes before and after differentiation.

Primary adipocytes and SVF were isolated from three different adipose tissues: gonadal WAT, subcutaneous WAT and Classical BAT. After collection and digestion, cells were filtered through a 250 μm mesh and left to stand on ice for 30 minutes. Primary adipocytes were collected (floating fraction), and the rest was further centrifuged (10 min, 700g). Remaining mature adipocytes were collected (floating fraction), and the SVF (pelleted fraction) was further processed by washing with DMEM media and centrifuging (10 min, 700g). Last, the supernatant was removed, and the pellet was processed for RNA analysis.

Immortalised brown and white adipocyte cell lines were generated by culturing the cells from the SVF of iBAT and scWAT of 10-week-old female 129Sv mice with retroviral-mediated expression of temperature-sensitive simian virus 40 (SV40) large T-antigen (H-2Kb-tsA58) (14).

HEK293, primary cultures and immortalised cell lines were cultured in DMEM/F12 medium (Sigma) supplemented with 10% FBS Serum 10500 (Gibco), 1% L-Glutamine and 1% Penicillin-Streptomycin antibiotics (Gibco). Mouse embryonic stem cells (mESCs) were cultured in Glasgow’s MEM (GMEM) (Invitrogen) supplemented with 10% FBS, 2 mM L-Glutamine (Sigma), 0.5% Penicillin-Streptomycin (Gibco), 0.1 mM MEM Non-Essential Amino Acids Solution (Gibco), 0.25% Sodium Bicarbonate solution (Gibco, 7.5% stock), 1 mM Sodium Pyruvate (Gibco), 0.1 mM 2-Mercaptoethanol (Gibco) and 1000 units/ml LIF (Merckmillipore) on 0.1% gelatin (Millipore)-coated flasks.

Immortalised BAT and WAT were cultured at 33°C and 5 % CO_2,_ whereas HEK293, primary cultures and mESCs were cultured at 37°C in a humidified incubator at 5% CO_2_. Cells were passaged when they reached 50 – 70% confluence.

### 5.2 Adipocyte Differentiation

Mouse brown and white adipocytes were initially seeded in plates pre-coated with 0.1% gelatin at 90% confluency one day prior to differentiation. When cells reached confluency, plates were transferred to the 37 °C incubator and differentiation was initiated. Two distinct protocols were employed to induce the differentiation of brown and white adipocytes. The components and their concentrations in the induction and maintenance media for white and brown adipocytes are detailed in Supplementary Tables 1 and 2, respectively. Human pluripotent stem cells were induced to differentiate into mature BAs, as previously described (15). mESCs were induced to differentiate into mature adipocytes following the established protocol described earlier (16).

### 5.3 RNA extraction and RT-qPCR

For total RNA isolation, cells were collected in Trizol reagent (Invitrogen) and isolated as described in the manufacturer’s protocol. In brief, cells were lysed in 1 ml TRIzol reagent. Chloroform (TRI reagent®: Chloroform 5:1 v/v) was added. Cell mixtures were vigorously shaken for 15 seconds, incubated at RT for 3 minutes and centrifuged at 12,000 x g for 15 minutes at 4 °C. Next, the upper aqueous phase was transferred to a new tube and twice the volume of isopropanol (Sigma-Aldrich) was added for precipitation. After washing with 75% ethanol, the pellets were air dried and dissolved in nuclease-free water. RNA quantity and quality were analysed by a Nano-Drop spectrophotometer (Thermo Scientific).

The extracted RNA was reverse transcribed into cDNA. Initially, the RNA samples were incubated with Amplification Grade DNase I (Sigma-Aldrich) for 15 minutes. To neutralize DNase activity, EDTA stop solution was added to each sample and incubated at 65°C for 10 minutes. For reverse transcription, M-MLV Reverse Transcriptase (Sigma-Aldrich) was utilised to generate complementary DNA strands. Each sample received the following components: 1 µl Random Hexamer Primer (Thermo Fisher), 1 µl dNTP Mix (Promega), and 4.5 µl water. After incubating for 10 minutes at 70 °C, the remaining components were added: 2 µl of 10x M-MLV Reverse Transcriptase Buffer, 1 µl M-MLV Reverse Transcriptase, 4.5 µl water, and 0.5 µl RNase OUT™ (Invitrogen). The reaction underwent incubation at 25°C for 10 minutes to facilitate the elongation of random primers. Subsequently, RNA was reverse transcribed at 37°C for 50 minutes, followed by denaturation at 80 °C for 10 minutes.

For relative quantification of miRNAs, the miRCURY LNA RT Kit (Qiagen) was used for reverse transcription into complementary DNA (cDNA). The concentration of DNase-treated cellular RNA was adjusted to 5ng/µl in RNase-free water and reverse transcribed according to the manufacturer’s protocol.

For quantitative real-time PCR (qRT-PCR), the Applied Biosystems QuantStudio 7 Pro real-time PCR system (Thermo Scientific) was used. qRT-PCR for genomic or protein-encoding genes was performed by utilizing SYBR® Green JumpStart™ Taq ReadyMix™ (Sigma-Aldrich) according to the manufacturer’s guidelines. The list of primer sequences is provided in Supplementary Table 3 and 4.

The qRT-PCR analysis of miRNA was carried out by using the miRCURY SYBR Green PCR Kit in combination with miRCURY LNA miRNA PCR primers (339306, QIAGEN) for mmu-miR-10b-5p (YP00205637), U6 snRNA (YP02119464), mmu-miR-709 (YP00205637), mmu-miR-706 (YP00205976), hsa-miR-138-5p (YP00206078), hsa-miR-196a-5p (YP00204386), hsa-miR-365a-3p (YP00204622), mmu-miR-455-3p (YP00205432), mmu-miR-222-3p (YP02119325) and mmu-miR-378a-3p (YP00204179) following the instructions of the manufacturer. The relative fold change was calculated using the formula 2^-ΔΔCt^.

### 5.4 Oil Red-O Staining

On the final day of adipocyte differentiation, cell monolayers were fixed with 4% paraformaldehyde in PBS for 15 minutes at room temperature. A 0.25% (w/v) Oil Red O stock solution was prepared in 60% isopropanol, then diluted 6:4 with water to create the working solution. The cells were stained with this solution for 1 hour. Excess stain was removed by briefly washing with 60% isopropanol, followed by PBS.

### 5.5 Luciferase assay and Mutagenesis assay

miR-10b targets were predicted with TargetScan (targetscan.org) and MirWalk (<u>mirwalk.umm.uni-heidelberg.de</u>). Mouse Gata6 (1045 bp) and Tubby (2128bp) 3′-UTRs were amplified by PCR (primers Supplementary Table 5) and cloned into the pRL-TK vector (Promega) using the XbaI and EagI restriction sites downstream of the Renilla reporter gene. Mutagenesis was performed using the QuickChange Site-directed Mutagenesis Kit (Agilent) according to the manufacturer’s protocol. The sequence accuracy of all clones was verified by sequencing.

For luciferase activity assays, HEK293 cells in 48-well plates were co-transfected with locked nucleic acid (LNA) miR-10b mimic (5 nM) or control oligos (Exiqon) with the luciferase reporters using lipofectamine RNAiMAX (Invitrogen). Luciferase activity was measured 48 h after transfection. The pGL3-Control vector (Promega) containing the firefly reporter gene was co-transfected into all cells to normalize the results. Firefly and Renilla luciferase activities were analysed using the Dual-Luciferase reporter assay system (Promega) according to the manufacturer’s instructions. Luciferase activity was quantified using a microplate reader, Infinite 200 Pro (Tecan). Renilla luciferase activity was normalized to Firefly luciferase activity.

### 5.6 sgRNA plasmid construction and introduction into mESCs

CRISPR sgRNA were designed using the Optimized CRISPR Design tool CHOPCHOP (17–19). Considering the significance of Drosha and Dicer processing sites during the process of miRNA biogenesis, the sgRNA sequence for miR-10b was designed within/adjacent to these sites (Figure 4.A). The sgRNA sequence for miR-10b was 5’-CCTGTAGAACCGAATTTGTGtgg-3’ (protospacer adjacent motif (PAM) sequences depicted in lower case letters). The sgRNA design was also evaluated for off-target effects using the web tool, CCTop - CRISPR/Cas9 target online predictor (20).

The chosen single-stranded oligonucleotides containing the guide sequence of the gRNA were annealed and cloned into the BbsI linearised px330-U6-Chimeric_BB_CBh-huSpCas9 vector (Addgene plasmid # 42230); (21). The non-targeting sgRNA (Addgene plasmid # 42230) was used as a negative control. The generated CRISPR-Cas9 vectors harbouring either the gRNA targeting miR-10b or a non-targeted control (NTC) were transfected into the cells (Supplementary Table 6). To generate clonal miR-10b KO cell lines, fluorescence-activated cell sorting (FACS) was utilized 48 hours post-transfection. Individual cells exhibiting GFP fluorescence were isolated through sorting using BD FACSMelody™ Cell Sorter (BD Biosciences, UK) and expanded. Cells were centrifuged to remove culture media, washed in DPBS with Ca^+2^ and Mg^+2^, and then suspended in DPBS prior to analysing and sorting them by FACS. Cells were initially gated for the intact cell population using forward scatter versus side scatter plots and then gated for transfected cells based on the presence of the transfected control. Transfected cells were gated for GFP-positive cells on the side scatter versus GFP plots. Following cell clone isolation and expansion, the clones were screened and identified by PCR, followed by High Resolution Melt Analysis (HRMA) (22). Sanger sequencing analysis of the miR-10b locus was used to confirm the indels created by CRISPR/Cas9. In addition, we generated three clonal non-targeted cell lines as a negative control by transfecting ESCs with the CRISPR/Cas9 plasmid harbouring NTC sgRNA.

### 5.7 Extraction of genomic DNA and High-Resolution Melt Analysis (HRMA)

DNA extraction: Half of the cells from the clonal cell colony were transferred to a new well for expansion, while the remaining cells were briefly centrifuged. To the pellet, 50 µL of QuickExtract™ DNA Extraction Solution (Cambio) was added and mixed thoroughly. The mixture was then subjected to a heat treatment at 65°C for 6 minutes, followed by 98°C for 2 minutes. Subsequently, the sample was centrifuged again to pellet the cell debris.

To screen TALEN-induced mutations by HRMA (23). A 124 bp fragment that included the entire genomic target site was PCR amplified. Primers flanking the target site were used to amplify the amplicon (Supplementary Table 6). PCR reactions contained 5 µL of SYBR® Green JumpStart™ Taq ReadyMix™ master mix (Sigma), 1 µL of each primer (10 µM), 2 µL of DNA and water up to 10 µL (nuclease-free water; Qiagen). HRMA data was collected on the Applied Biosystems QuantStudio 7 Pro real-time PCR system (Thermo Scientific) and analysed using Excel (Microsoft Office).

### 5.8 Transfection (miRNA inhibitors, miRNA mimics and siRNA)

All miRNA mimics and miRNA inhibitors were purchased from QIAGEN. All oligonucleotide sequences and catalogue numbers are shown in Supplementary Table 7. RNAiMAX (Invitrogen) was used to deliver miRNA inhibitors and mimics into cells as per the manufacturer’s instructions. Silencer® Select siRNA against Tub (assay ID: s75577, cat. 4390771) and Silencer™ Select Negative Control No. 1 siRNA (cat. 4390843) were purchased from Thermo Fisher Scientific. Briefly, BAs or WAs were plated in twelve-well dishes at densities ranging between 200 and 250 K cells/well in media without antibiotics and transfected with either 50 nM miRNA inhibitors or 5 nM miRNA mimics. For Tub knockdown assays, cells were treated with 5 nM of siRNA. Forty-eight hours later, cells were treated with induction medium. Cells were also treated with a negative control that did not exhibit any homology to the mouse sequence in the NCBI and miRBase databases.

### 5.9 Measurement of Oxygen Consumption Rates

Oxygen consumption rates (OCR) were measured using the Seahorse XFe24 Analyzer (Seahorse Bioscience). IBAT and scWAT cells were seeded into 0.1% gelatin-coated 24-well Seahorse Microplates (Seahorse Bioscience) at a density of 10,000 cells/well and treated with miR-10b inhibitor or mimic. The following day, cells were induced to differentiate using the standard protocols outlined in Supplementary Tables 1 and 2. Each experimental group included at least five replicates. The assay was conducted in sterile, unbuffered Assay Media prepared with Seahorse base media (Seahorse Bioscience) at 37°C (pH 7.4), and supplemented with Sodium Pyruvate (1mM, Sigma), Glucose (10 mM, Sigma). Mature adipocytes were washed twice in this supplemented seahorse media to fully remove the maintenance media and were then incubated for 1 hour at 37°C with no CO_2_. Mitochondrial function and potential were analysed by sequential injection of 1 µM oligomycin (ATP synthase inhibitor), 1 µM Carbonyl cyanide-4 (trifluoromethoxy) phenylhydrazone (FCCP, mitochondrial uncoupler) together with 2 mM pyruvate to achieve maximal respiration, and 2 µM rotenone and antimycin A (mitochondrial complex I and III inhibitors, respectively). Seahorse results were normalised to cell count using Hoechst 33258 fluorescent dye signal quantified immediately after Seahorse analysis (24). Cells were counted at the end of the experiments to normalize the OCR, ECAR, and PER readings. Respiration, acidification, and ATP production rates were calculated following the Agilent Seahorse XF Technology instructions.

### 5.10 Western Blotting

Whole-cell lysates for Western blotting were extracted with RIPA lysis buffer containing Halt™ Protease Inhibitor Cocktail (ThermoFisher Scientific). Equal amounts of protein were resolved on 10% SDS-PAGE gels and transferred onto nitrocellulose membrane by using a Trans-Blot Turbo System (BioRad) set at 1.3 A/25 V/10 min. Membranes were blocked with 3% non-fat milk for 1 h at 20 °C and incubated overnight at 4 °C with the following primary antibodies (1:1000): UCP1 (ab10983, Abcam), HSP60 (15282-1-AP, Proteintech), TUB (17928-1-AP, Proteintech) and GAPDH (MAB374, Sigma Aldrich). Membranes were then incubated for 1 hour at room temperature with the appropriate secondary antibody Goat Anti-Rabbit IgG (H+L) or Goat Anti-Mouse IgG (H+L), conjugated to IRDye® 800CW or IRDye® 680RD (926-32211, 926-68071, 926-68070, 926-32210, Licor Biosciences) at a 1:15,000 dilution in TBS/T containing 0.1% Tween. Washing steps were repeated after the incubation. Membranes were visualised with Odyssey CLx Imager (Licor Biosciences). Densitometry was performed via EmpiriaStudio (Licor Biosciences) software to measure the expression of the protein of interest.

### 5.11 RNA-sequencing analysis

For small RNA sequencing, a total amount of 3 μg total RNA per sample was used as input material for the small RNA library. Sequencing libraries were generated using NEBNext® Multiplex Small RNA Library Prep Set for Illumina® (NEB, USA). The clustering of the index-coded samples was performed on a cBot Cluster Generation System using TruSeq SR Cluster Kit v3-cBot-HS (Illumina) according to the manufacturer’s instructions. After cluster generation, the library preparations were sequenced on an Illumina platform and 50bp single-end reads were generated. Small RNA analysis was performed by Novogene Ltd (Cambridge, UK). Briefly, Raw FASTQ reads were processed using custom Perl and Python scripts to remove reads containing ploy-N, with 5’ adapter contaminants, without 3’ adapter or the insert tag, containing ploy A or T or G or C and low-quality reads from raw data. All the downstream analyses were based on the clean data with high quality. The small RNA tags were mapped to reference sequence by Bowtie (25) without mismatch to analyse their expression and distribution on the reference. Mapped small RNA tags were used to looking for known miRNA. miRBase20.0 was used as reference. miRNA expression levels were quantified using TPM (transcripts per million) based on the criteria outlined by Zhou et al. (26). The normalization formula used was: Normalized expression = (mapped read count / total reads) × 1,000,000. The miRNA expression dataset can be accessed through ArrayExpress under accession number E-MTAB-15230.

For bulk RNA sequencing, a total amount of 400 ng total RNA per sample was used as input material for the RNA library reparation of the RNA library and transcriptome sequencing was conducted by Novogene Ltd (Cambridge, UK). Regarding the transcriptomic analysis performed on mESCs differentiated to mature adipocytes, independent biological sample replicates from three different NTC clones and KO clones (Supplementary Table 9) were used for RNA sequencing. For the transcriptomic analysis performed on mouse BAT treated with miR-10b inhibitor or negative control, 4 independent biological sample replicates were used for RNA sequencing. Library Construction and Sequencing and conducted by Novogene Ltd (Cambridge, UK). RNA sequencing data of miR-10b +/+ and miR-10b –/– mESCs during adipocyte differentiation (Days 0, 12, and 27) are available under ArrayExpress accession E-MTAB-15151. Data from the RNA sequencing analysis of miR-10b-5p knockdown during brown adipogenesis will be available under E-MTAB-15152.

Bioinformatic analysis of the RNA sequencing samples was performed in the R statistical computing environment version 4.3.2 and RStudio version 2024.12.0 using a published pipeline adopted from an open-source toolkit (27).

Kallisto (28) was used to generate read counts from a transcriptome reference. Kallisto transcript abundance measurements were imported into RStudio using the R package tximport (version 1.28.0) (29) for filtering and normalization. The Bioconductor/R package edgeR *cpm* function was used to create a list of counts per million (cpm) per transcript (30). Genes with fewer than 10 cpm were filtered out, retaining only those with read counts exceeding 10 in at least 3. Genes included in the analysis had >1 cpm in at least 3 samples. The edgeR (version 4.0.4) function *calcNormFactors* was used to normalize the filtered data using the TMM method (Trimmed Mean of M-values) to the data set (31). The voom function from the Bioconductor/R package *limma* (version 3.58.1) (32) was used to establish a mean-variance relationship and model its trend. The lmFit function was then applied to fit a linear model to the data, while makeContrasts was utilized to define contrasts in the design matrix. Bayesian statistics were applied to the linear model fit using the eBayes function. To identify differentially expressed genes (DEGs), the decideTests function was used, applying multiple testing correction via the Benjamini-Hochberg method and strict cutoff criteria (corrected p-value ≤ 0.05 and log₂ fold change ≥ 1 to generate a list of significantly regulated DEGs (33). Principal Component Analysis (PCA) plots were generated by using the built-in R functions prcomp(). The Bioconductor/R packages *ggplot2* (version 3.4.4) was used to create volcano plots. Heatmaps were generated using the heatmap.2 function of the R package *ggplot2*. The Bioconductor/R packages *msigdbr* (version 7.5.1), *clusterProfiler* (version 4.10.0) and *enrichplot* (version 1.22.0) were utilized to conduct gene set enrichment analyses using the C2 curated database from the Molecular Signatures Database (MSigDB). The Bioconductor/R package gprofiler2 (version 0.2.2) (34) was used to perform GO pathway analysis.

### 5.12 Statistical Analysis

Experimental data are presented as mean ± standard error of the mean (SEM) and were analysed with Prism software (Graph pad, Inc.). The normal distribution of the data was assessed by a Shapiro-Wilk normality test. For normally distributed data, group comparisons were performed using two-tailed t-tests, one-way ANOVA, or two-way ANOVA, with Sidak’s post hoc test for multiple comparisons. For data that were not normally distributed, the Kruskal–Wallis test was used, followed by Dunn’s multiple comparison test. Details of the specific tests used in each experiment are provided in the corresponding figure legends. A significance level of *p* < 0.05 was deemed significant.

### 5.13 Cellular Senescence

Cell senescence was visualized via a β-galactosidase staining protocol (35). For positive control, we treated cells with 500 µM H_2_O_2_ prior to β-galactosidase staining. Stained cells, which appear as light blue cells, were observed via light microscopy at ×20 magnification using a DMi1 Inverted Microscope, (Leica Microsystems).

### 5.14 miRNA PCR Array

We determined differentially secreted miRNA expression in conditioned media from undifferentiated and differentiated brown and white adipocytes conditionally immortalised cell lines using miRCURY LNA^TM^ Universal RT microRNA PCR Mouse&Rat panel I+II comprehensive for 752 miRNA assays. Cells were initially seeded in a 6 well plate and left in serum-free media for 24 hours to allow miRNA accumulation. Media samples were collected and processed through serial centrifugations, filtered, and then stored at −80 °C. Subsequent steps were carried out at Exiqon Service, where RNA was isolated and reverse transcribed. The resulting cDNA was analysed using the miRCURY LNA™ Universal RT microRNA PCR Mouse & Rat panel I+II. The scanning was performed using the Agilent G2565BA Microarray Scanner System (Agilent Technologies, Inc., USA) and the image analysis was carried out through the ImaGeneR 9 (miRCURY LNATM microRNA Array Analysis Software, Exiqon, Denmark).

### 5.15 Immunofluorescence

Cells plated on 0.1% gelatin-coated coverslips, washed in PBS and fixed with 4% paraformaldehyde for 10 min and permeabilized for 5 min with 0.1% Triton X-100 in PBS. Cells were washed with PBS and blocked with 5% goat serum in PBS for 1 h at RT, followed by overnight incubation with Tub antibody (1:400; 17928-1-AP, Proteintech) diluted in 1% BSA in PBS. Cells were washed with PBS and incubated for 1 h at RT with goat anti-rabbit IgG Alexa Fluor 488 (Invitrogen). Cells were rinsed with PBS and incubated in LipidTOX-Deep Red (1:2000, Thermo-Fisher) for 1 h at RT. Following incubation, cells were stained with 300 nM DAPI for 40 min and coverslips were mounted on glass slides with ProLong antifade reagent. Images were obtained with either a Leica SP5 confocal microscope or Nikon AX R Confocal Microscope. Images were processed using FIJI/ImageJ software.

### 5.16 Cell proliferation assay

Cells were seeded at 1 million per well in a 6-well plate, incubated for 48 hours, then counted and passaged. This process was repeated every 48 hours for 8 passages, with reseeding at 1 million cells per well each time. Cell counts were recorded at each passage to monitor proliferation trends.

## 6 Results

### 6.1 Identification of differentially expressed miRNAs between WAT and BAT

To identify miRNAs differentially expressed among undifferentiated and differentiated brown and white adipocytes, we profiled miRNA expression of preadipocytes isolated from the SVF of scWAT and iBAT, undifferentiated or differentiated into mature brown and white adipocytes by Small RNA Sequencing. Principal component analysis (PCA) of cellular miRNAs showed that samples clustered into 4 distinct groups, indicating that each group had a distinct miRNA profile (Figure 1A). Small RNAseq analysis identified 126 differentially expressed miRNAs in mature adipocytes and pre-adipocytes based on log_2_ fold change of 2 and adjusted p-values <0.05 with Benjamin-Hochberg correction (Figure 1B, C). We focused on brown upregulated miRNAs to identify the miRNAs that might play a critical role in the brown phenotype of adipocytes. In differentiated BAT versus WAT, there were 28 upregulated and 42 downregulated miRNAs, whereas in BAT versus WAT preadipocytes, there were 61 upregulated and 45 downregulated miRNAs based on log2 fold change of 1 and presented a p value ≤0.05 (Figure 1D). Among significantly regulated miRNAs, miR-10b was identified as consistently enriched in brown adipocytes (Figure 1C, D, E).

**Figure 1.**
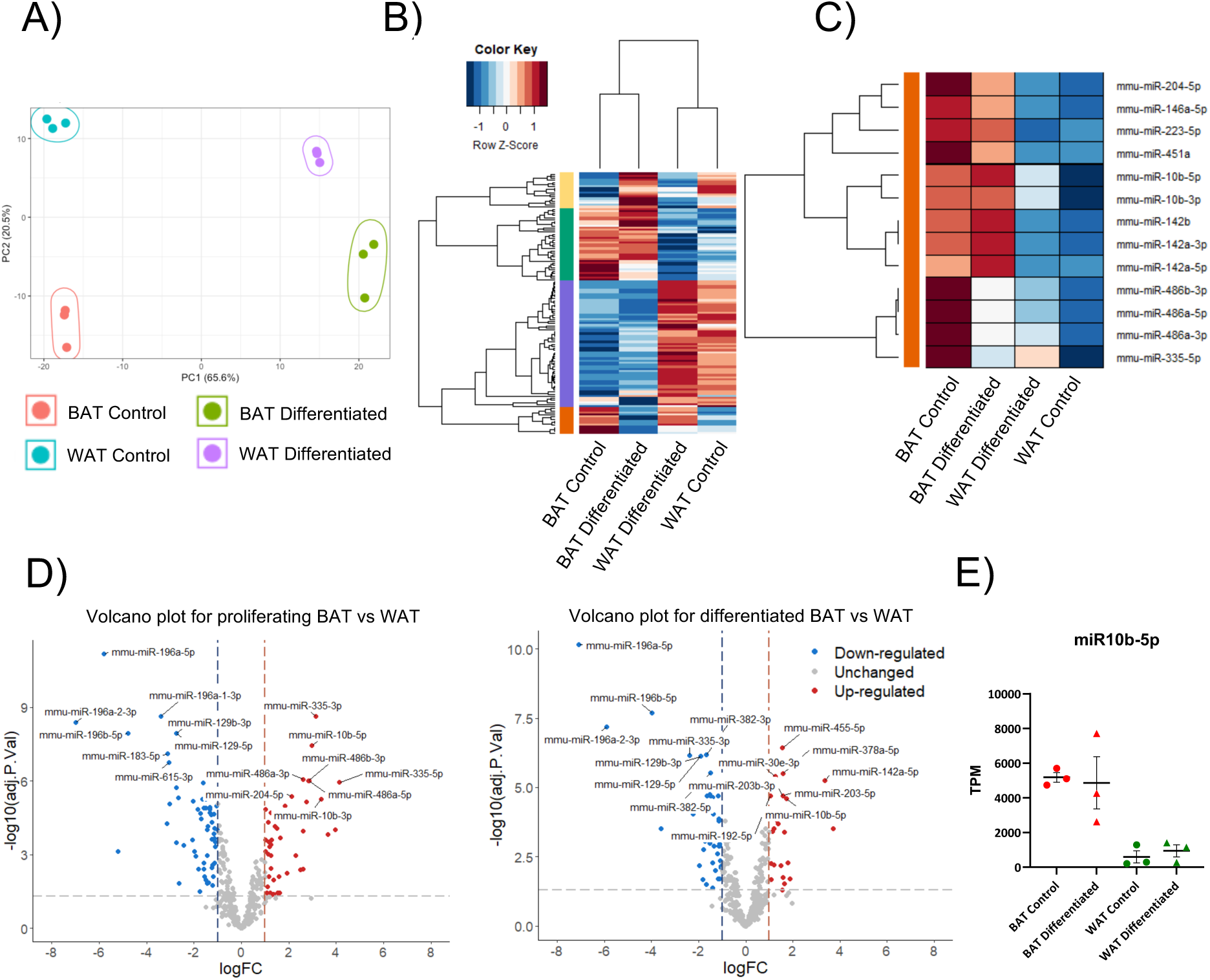
Small RNA sequencing identifies differentially expressed miRNAs during brown and white preadipocyte differentiation to adipocytes. A) PCA plot. B) The most dynamically expressed miRNAs are shown in the heatmap. C) The right heat map includes a number of selected miRNAs that are highly expressed in BAT compared to WAT (orange module). D) Volcano plots for the comparison between undifferentiated and differentiated BAT and WAT primary cultures. E) Scatterplot represents the expression levels of miR-10b-5p as measure in TPMs. Solid horizontal black line represents the mean ± SEM. TPM = transcripts per million.

### 6.2 Profile of differentially secreted miRNAs among undifferentiated and differentiated WAT and BAT

miRNAs have also been found extracellularly in biological fluids. Circulating miRNAs may facilitate paracrine and endocrine crosstalk between tissues and cells, thereby influencing gene expression and cellular activities. Adipose tissue is a major contributor to the pool of secreted miRNAs. Consequently, changes in adipose tissue mass or function, common in various metabolic conditions, can alter circulating miRNAs, which then exert systemic effects (36). We analysed the differentially secreted miRNAs in conditioned media from undifferentiated and differentiated brown and white adipocytes. We identified 143 miRNAs common to all samples, with an average of 225 miRNAs detectable per sample. Variation among the assays was analysed by PCA, unsupervised two-way hierarchical clustering and pairwise volcano plots (Figure 2).

**Figure 2.**
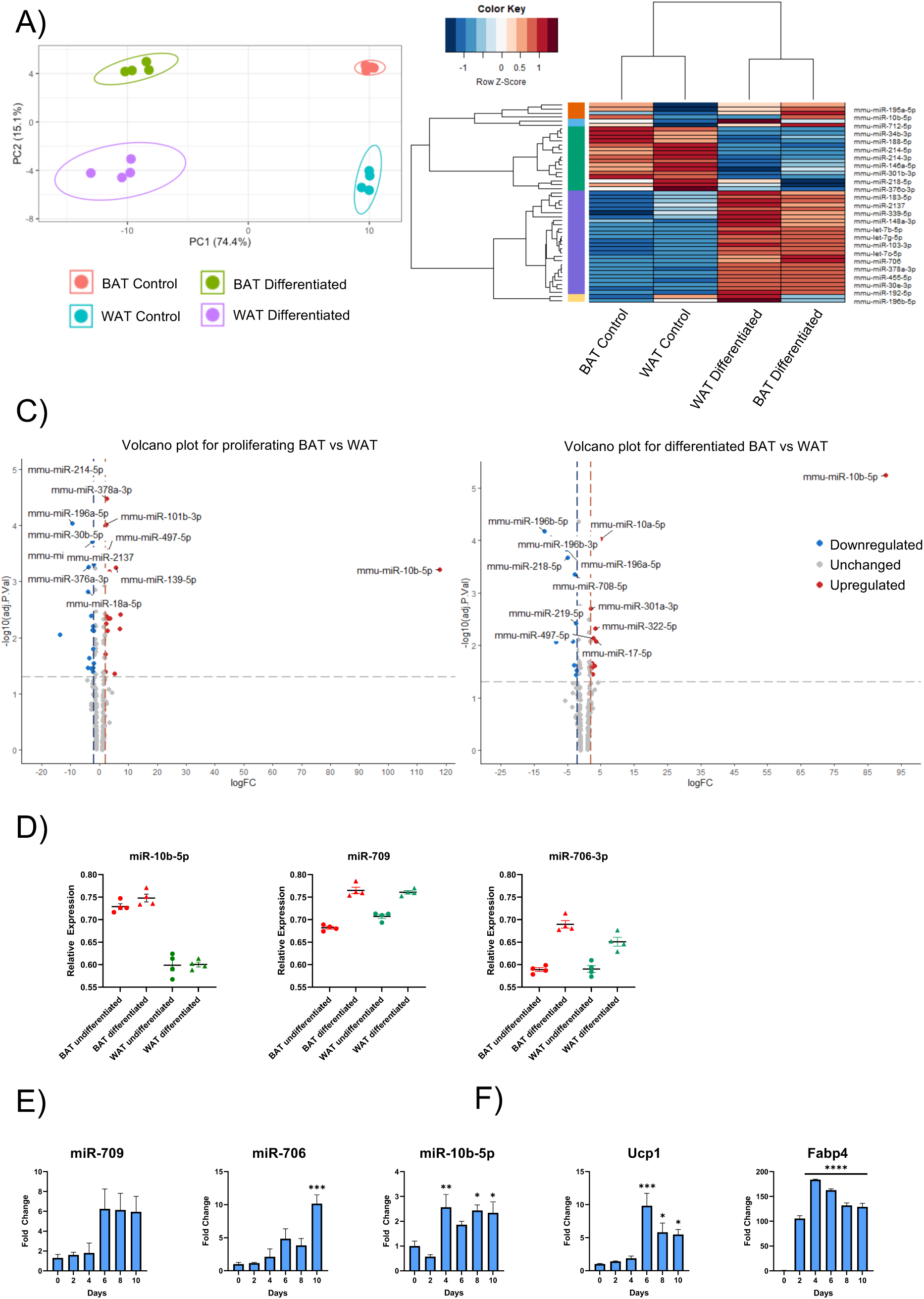
Identification of differentially secreted miRNAs during Brown and White preadipocyte differentiation into mature adipocytes. A) PCA and B) Heat Map and Unsupervised Hierarchical Clustering were conducted on the 50 miRNAs with the highest coefficient of variation based on normalized (dCq) values across all samples. C) Volcano plots showing the relationship between the p-values and the ddCq during the pairwise comparison of undifferentiated BAT vs WAT and differentiated BAT vs WAT. Highlighted spots are microRNAs exhibiting a log2 fold change of 2 or above and with p-values below 0.05 after Benjamini-Hochberg correction for multiple testing. The top 6 miRNAs are labelled. D) Scatterplots represent the relative secretion levels of miR-10b-5p, miR-706 and miR-709. Data normalized with UniSp3. E) Expression profiles of miR-10b-5p, miR-706 and miR-709 during BAT differentiation. F) Expression levels of Ucp1 and Fabp4 during BAT differentiation. Data were analysed using one-way ANOVA. Each day’s mean is compared to the mean of day 0, with significance assessed for each comparison. Data represent the mean ± SEM (*p< 0.05, **p< 0.005, ***p< 0.0005, ****p < 0.0001).

The PCA plot shows clearly that the four sample groups have very distinct miRNA profiles. The most significant contributor to the observed variance was the differentiation status of the cells (Figure 2A). Further analysis with heat map and two-way hierarchical clustering showed two distinct signatures, one for the tissue type (BAT versus WAT) and one for the differentiation state between BAT and WAT (Figure 2B). Overall, several secreted miRNAs were found to be upregulated during brown differentiation when compared to WAT differentiation (Figure 2C). The most prominent secreted miRNA that exhibited the highest log fold change during the pairwise comparison of undifferentiated BAT vs WAT and differentiated BAT vs WAT was miR-10b-5p (Figure 2C).

Furthermore, miR-706 and miR-709 were also identified amongst the significantly differentially secreted miRNAs during differentiation of BAT and WAT (Figure 2D). Thus, we identified three miRNAs (miR-10b, miR-706 and miR-709) as potentially involved in brown and white adipogenesis. Validation of these miRNAs identified from miRNA profiling arrays and RNA sequencing was performed by quantitative real-time PCR using locked nucleic acid (LNA) detection primers. An upward trend in miR-709 expression was observed from day 6, whereas miR-10b and miR-706 were significantly upregulated during a BAT differentiation time course (Figure 2E). As an assessment of BAT differentiation, we also analysed the expression of Fabp4 and Ucp1 (Figure 2F). As expected, we observed significantly higher expression of key thermogenesis marker Ucp1 and adipogenesis marker Fabp4 during the later stages of BAT differentiation.

### 6.3 miR-10b is upregulated during ESCs differentiation to mature adipocytes

#### 6.3.1 Mouse ESCs

A large part of our knowledge of the molecular pathways governing adipogenesis has been acquired using cellular model systems such as immortalized murine cell lines (3T3-L1, 3T3-F442A, ob1771 and OP9), multipotent murine embryonic cell line C3H10T1/2, and primary culture models (SVF of fat depots) (37). These models primarily focus on the late stages of the differentiation program. Thus, the sequence of events involved in the commitment of adipose tissue stem cells/precursors to preadipocytes remains largely unexplored. To study the involvement of the chosen miRNAs during the early stages of preadipocyte commitment, we exposed mESCs to an optimised protocol (16) to induce the differentiation of mESCs to terminally differentiated adipocytes (Figure 3A). QPCR analysis at various time points during differentiation confirmed a significant upregulation of adipocyte markers Ucp1 and Fabp4 (Figure 3B), along with miRNAs that have previously shown to regulate adipogenesis, including miR-138-5p (38,39), miR-196a-5p (40), miR-365-3p (41,42), miR-455-3p (43), miR-222-3p (44) and miR-378a-3p (45–47) (Figure 3C). miR-10b-5p showed a dynamic expression profile during the time course (Figure 3D). A nearly 389-fold increase of miR-10b was observed during pre-adipocyte development at day 12 compared to day 0. The expression of miR-706 and miR-709 did not appear to alter significantly during ES differentiation (Figure 3D). We thus hypothesized that induction of miR-10b-5p is a critical event for adipocyte development.

**Figure 3.**
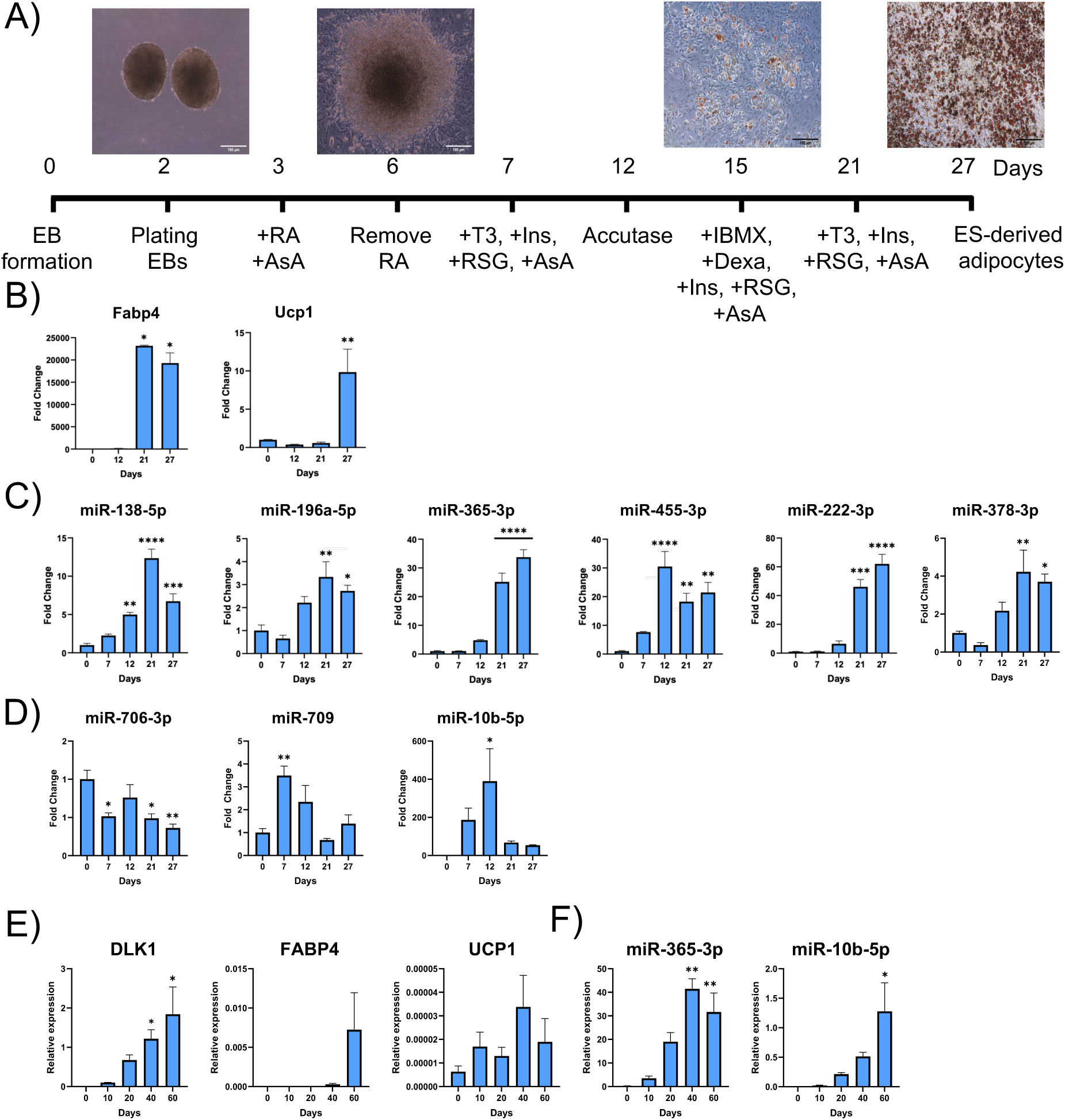
Adipogenesis markers and miRNA expression during ESCs differentiation to mature adipocytes. A) Stepwise adipocyte differentiation protocol from mouse embryonic stem cells. Protocol adapted from (Cuaranta-Monroy I. et al. 2014). B) Fabp4 and Ucp1 expression were elevated during ESCs differentiation. C) qPCR analysis of miRNAs previously shown to regulate adipogenesis. D) The expression of miR-706, miR-709 and miR-10b was determined by SYBR qRT-PCR. E) Expression levels of DLK1, FABP4, and UCP1, and F) miR-365 and miR10b-5p, monitored during the differentiation process of hPSCs into BAs. mRNA and miRNA expression were normalised to the housekeeping genes L19 and U6, respectively. Data from FABP4 (BA), miR-222 (BA), miR-709 (BA) and DLK1 (hESCs) analyses were evaluated using the Kruskal-Wallis test, while the remaining gene datasets were analysed using one-way ANOVA. Each day’s mean is compared to the mean of day 0, with significance assessed for each comparison. Data represent the mean ± SEM (*p< 0.05, **p< 0.005, ***p< 0.0005, ****p < 0.0001).

#### 6.3.2 Human ESCs

The stepwise differentiation and functional characterization of paraxial mesoderm cultures towards an adipogenic fate has recently been modelled *in vitro* using hESCs (48). To study the expression of miR-10b in human adipocyte differentiation, human pluripotent stem cells (hPSCs) were differentiated into BAs. RNA samples were collected at different time points for qRT-PCR analysis. Induced pluripotent stem cell (iPSC) differentiation to fully mature BAs was confirmed by the upregulation of preadipocyte and brown adipogenic markers, including DLK1, FABP4 and UCP1 (Figure 3E). BA development was also corroborated by the upregulation of a known adipogenic miR-365-3p (41,42) (Figure 3F). In addition, miR-10b-5p expression steadily increased during iPSC differentiation (Figure 3F), supporting that miR-10b might regulate human BA differentiation.

Taken together, miR-10b is expressed and regulated across various cellular models of brown adipogenesis in mice and humans. These findings suggest that miR-10b may be essential not only to the formation and differentiation of brown adipose tissue but also to the overall regulation of metabolic processes.

### 6.4 CRISPR/CAS9 knockout of miR-10b

Because miR-10b is potentially relevant to the development of mature adipocytes, we sought to determine its specific role during adipogenesis by generating its functional knockout (KO) in E14 mESCs using CRISPR/Cas9 engineering technology. In the mouse genome, miR-10b resides in the intronic region of two overlapping genes, Hoxd3 and Hoxd4 (Figure 4A). Given the significance of Drosha and Dicer processing sites during the canonical pathway of miRNA biogenesis (49), we constructed the CRISPR/Cas9 vector to contain a single guide (sgRNA) designed to target within/adjacent to these sites in miR-10b. We identified knockout events in 15% of colonies (6 out of 40). Sanger sequencing confirmed CRISPR/Cas9-induced deletions and indels in four clones (Figure 4B). Sequence alignment showed complete disruption of the terminal loop, Drosha, and Dicer sites in three KO clones (3G6, 1F10, 3E6), while the fourth clone (3F4) had only the Drosha site affected.

**Figure 4.**
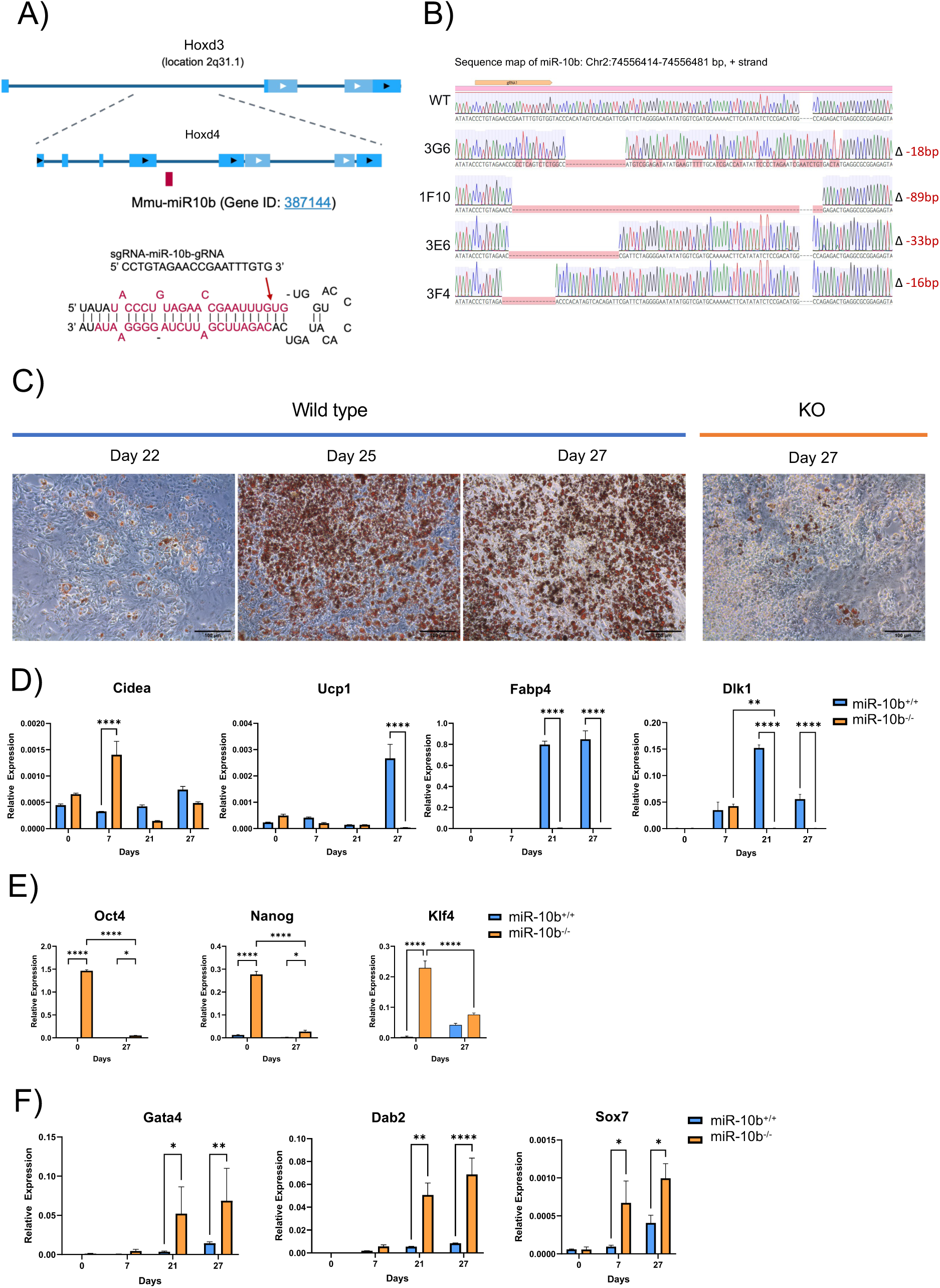
The loss of miR-10b significantly compromised adipogenesis. A) Design of sgRNAs flanking the miR-10b precursor sequence located in the intronic region of Hoxd3 and Hoxd4. B) Detection and confirmation of indels generated by CRISPR/Cas9 editing. DNA sequencing confirms the deletions (Red) by CRISPR/Cas9. C) The degree of differentiation was evaluated by visualising the amount of lipid accumulation on days 22, 25 and 27 of differentiation. A representative bright-field image from wild-type and miR-10b KO cultures depicting the progress of differentiation. Lipid droplets were stained with Oil-Red-O. Scale bar represents 100 µm. D) mRNA expression of the adipose-specific markers Cidea, Ucp1, Fabp4 and preadipocyte marker Dlk1 was measured by qRT-PCR. E) Stem cell markers OCT4, Nanog and KLF4, F) specific endoderm markers, Gata4, Dab2 and Sox7 were upregulated in KO clones. Data (n ≥ 3) are shown as means + SEM (two-way ANOVA test, *p< 0.05, **p< 0.005, ***p< 0.0005, ****p < 0.0001).

Using CCTop - CRISPR/Cas9 target online predictor (20,50), we predicted the off-targets of miR-10b sgRNA (Supplementary Table 8). Most targets had four base mismatches in the seed region and were in exonic, intronic, and intergenic regions. We did not analyse gene expression because the targets were primarily intronic or intergenic, or due to previous research findings that the presence of four or more base mismatches eliminates the detectable SpCas9 cleavage in most loci (51,52).

We compared doubling times of WT and miR-10b KO cells to assess the impact on proliferation rates and found no significant difference (Supplementary Figure S1A). Alkaline Phosphatase (AP) activity was used to evaluate self-renewal capacity in the presence and absence of LIF. Both KO and WT cells formed more undifferentiated colonies with LIF, but KO cells, derived from a clonal population, showed a higher number of undifferentiated colonies than WT (Supplementary Figure S1B).

### 6.5 Knockout of miR-10b severely compromises adipocyte differentiation

Loss of miR-10b in all four KO clones significantly impaired the differentiation process into mature adipocytes. In depth analysis was performed on KO clone 1F10. There was a clear reduction of lipid accumulation as judged by Oil Red O staining in KO compared to WT cells at day 27 of the differentiation treatment (Figure 4C). Expression of Fabp4 and Ucp1 was profoundly reduced in KO compared to WT cells at day 27, further supporting that adipogenesis was impaired in the absence of miR-10b (Figure 4D). It is noteworthy that in the pluripotent state, there was elevated Ucp1 expression in miR-10b KO compared to WT cells that was lost during the commitment and differentiation process (Figure 4D).

Remarkably, the transcript abundance of Preadipocyte factor 1 (Pref-1/Dlk1), a molecular gatekeeper of adipogenesis (53), was consistently higher in WT compared to KO at 21 and 27 days of differentiation (Figure 4D). Furthermore, Dlk1 expression was elevated only at day 7 in the KO and then diminished, suggesting that the absence of miR-10b expression hindered the commitment stages and prevented the cells from differentiating into mature adipocytes. The levels of Cidea remained largely unchanged throughout the differentiation process, apart from day 7, where KO cells exhibited significantly elevated expression levels compared to WT cells (Figure 4D).

Cellular senescence is a key contributor to impaired adipogenesis, leading to dysfunctional adipose tissue formation and function (54,55). To investigate whether cells entered senescence due to depleted miR-10b, we stained mature adipocytes for β-galactosidase and observed them at ×20 magnification. Microscopic examination revealed that KO cells did not display signs of senescence, as indicated by the absence of cells staining positive for β-galactosidase activity (Supplementary Figure S1C).

In agreement with the commitment steps being hindered in the absence of miR-10b, the expression levels of three major pluripotent genes, Nanog, Oct4, and Klf4, were highly elevated in KO cells at Day 0 and Day 27 compared to WT cells (Figure 4E). Moreover, expression of endoderm markers (Gata4, Dab2, and Sox7) was enriched in KO cells compared to WT cells (Figure 4F).

### 6.6 Impact of miR-10b Depletion on Stem Cell Commitment and Lineage Determination

To uncover the pathways influenced by miR-10b depletion, a transcriptomic analysis was conducted on samples collected on days 0, 12, and 27 (Supplementary Table 9). The PCA plot shows no clear segregation between the wild type and KO clones at day 0 (Supplementary Figure S2). At Day 12, there is a clear separation of wild-type and miR-10b KO samples, with further separation occurring by day 27 of differentiation, suggesting that the gene expression profiles of the two groups are distinctly different (Supplementary Figure S2). The main factor affecting the variance we observe appears to be the differentiation status of the cells. Transcriptomic analysis revealed 2304 genes with significant differential expression during key stages of mESC differentiation to mature adipocytes across both phenotypes (log_2_ fold change >2, adjusted p-values ≤0.05) (Figure 5A). To assess the differentially expressed genes (DEGs) between the two groups through days 0, 12 and 27 of adipogenesis, we utilized criteria where the adjusted p-value was ≤ 0.05 and the absolute value of log2 fold change ≥ 1.

**Figure 5.**
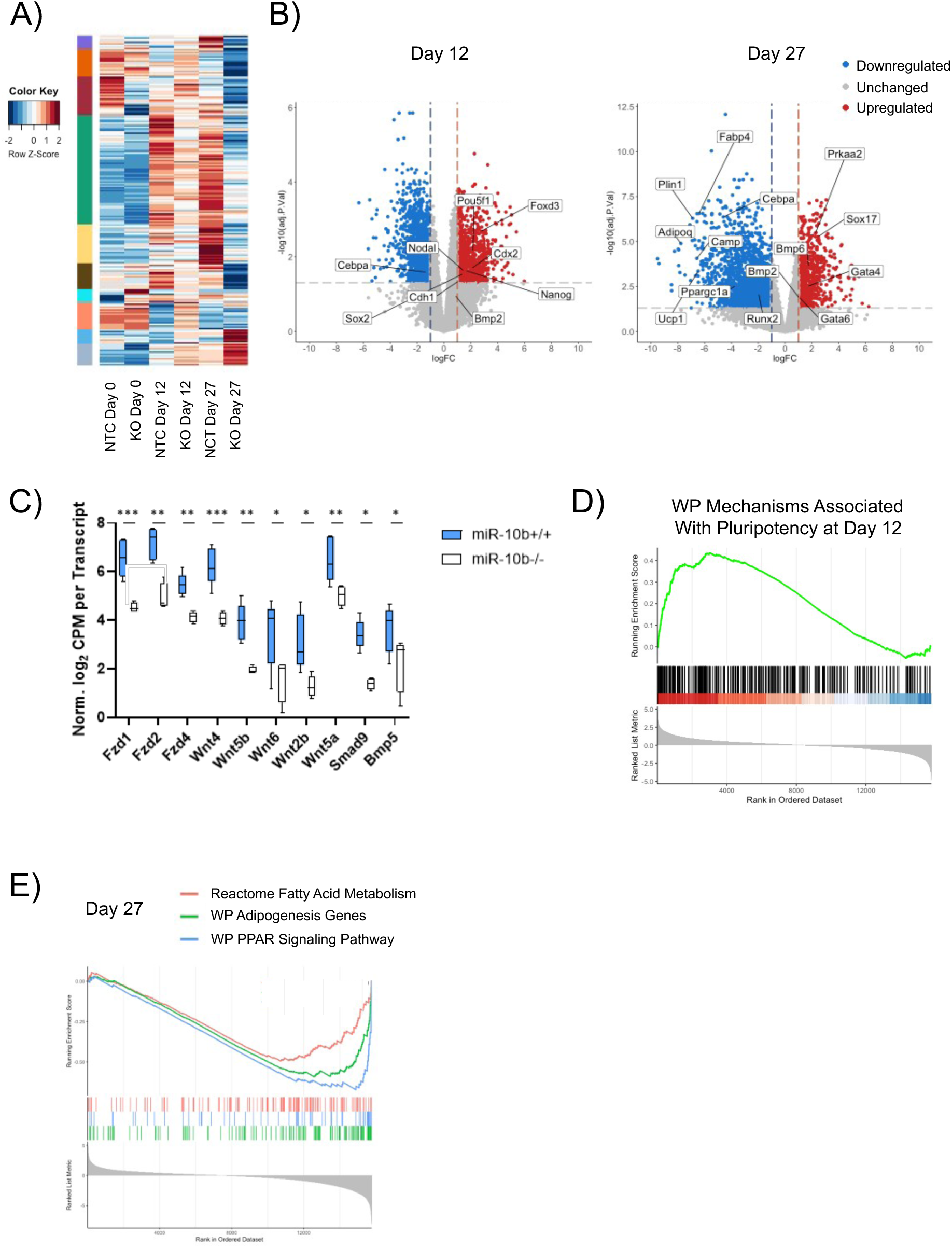
Transcriptomic analysis of mESCs either with or without depletion of miR-10b, during differentiation into mature adipocytes. A) The Heatmap displays the results of hierarchical cluster analysis applied to 2304 commonly observed DEGS between the different cell phenotypes during the time points of day 0, 12, and 27. These DEGs were identified based on a log2 fold change greater than 2 and adjusted p-values exceeding 0.05. B) Volcano plots depicting the expression profile of genes in mESCs treated with gRNA targeting miR-10b or NTC vector and differentiated at Day 12 and 27 (log2(FC) > 1 and p-adjusted value > 0.05). Negative logFC (red) = upregulated in KO; positive logFC (blue) = upregulated in mESCs treated with NTC vector. C) Normalized and log₂-transformed cpm for each transcript of Frizzled receptor family members and Wnt family genes, illustrating transcriptional activity in response to miR-10b depletion. Data were normalized using the TMM (Trimmed Mean of M-values) method via the calcNormFactors function in edgeR (v4.0.4). Adjusted p-values were calculated using the Benjamini–Hochberg method (p < 0.05, p < 0.005, p < 0.0005, p < 0.0001). Centre lines represent the medians; whiskers extend to the minimum and maximum values. D) GSEA at day 12 indicated a striking enrichment of transcripts involved in the mechanisms associated with pluripotency, whereas E) GSEA results from day 27 dataset exhibited a suppression in the levels of transcripts associated with Fatty Acid Metabolism, Adipogenesis Genes and PPAR Signalling Pathway in the KO group.

In total, 499 genes showed increased expression, while 679 genes exhibited decreased expression at day 0 (Supplementary Figure S2B). At day 12, during the commitment stage, 898 genes were upregulated, and 1147 genes were downregulated. On day 27, during the late phase of differentiation, 820 genes were upregulated, and 1981 genes were downregulated (Figure 5B).

Gene set enrichment analysis (GSEA) of the day 12 dataset demonstrated a striking enrichment of transcripts associated with mechanisms related to pluripotency (Figure 5D). Notably, Nanog, a homeodomain protein that is transcribed exclusively in pluripotent cells within ESCs, along with its transcription activators Pou5f1 (Oct4), Sox2 and Foxd3 (56) were upregulated in the miR-10b KO cells compared to control samples (Figure 5B). Moreover, various genes encoding members of the Frizzled receptor family (Fzd2, Fzd1, and Fzd4), which are involved in the WNT signalling pathway as well as members of the Wnt family (Wnt4, Wnt5b, Wnt6, Wnt2b, and Wnt5a) were shown to be downregulated in the KO group. Suppression of these proteins may decrease the activation of the WNT pathway (57,58), leading to a possible alteration in cell fate decisions and favouring the endodermal lineage (Figure 5C). Furthermore, genes involved in bone morphogenetic protein (BMP) signalling such as Smad9 and Bmp5 (59,60), were also downregulated in the KO group, which could lead to reduction of BMP signalling (Figure 5C), which is crucial for mesodermal commitment (61). These observations are consistent with the results indicating an elevation in the expression of specific markers associated with the endodermal lineage during adipogenesis treatment of KO mESCs (Figure 4F).

At the later stages of differentiation (day 27), transcriptomic analysis revealed an increase in the expression of adipogenic markers, including Fabp4, Plin1, Adipoq, Cebpa, Ucp1, Ppargc1a, and Runx2 in the wild-type clones (Figure 5B). This collective upregulation supports various aspects of adipocyte differentiation, lipid metabolism, and adipose tissue function, thereby promoting the adipogenic process. Consistent with the finding of suppressed terminal stem cell differentiation in miR-10b KO cells, GSEA at day 27 revealed that key signalling pathways associated with fatty acid metabolism, adipogenesis genes and PPAR signalling pathway were suppressed in the KO group (Figure 5E). Interestingly, the preference for the endodermal lineage observed in the KO group at day 12 persisted until day 27, as evidenced by the increased expression levels of Sox17, Bmp2, and Gata6 (Figure 5B). Taken together, these findings demonstrate that the loss of miR-10b appears to play a role in cell fate determination during embryonic development, leading to reduced mesodermal commitment and a preference for endodermal commitment.

### 6.7 Gata6 is a direct target of miR-10b

TargetScan was utilized to predict potential targets of miR-10b to investigate the molecular process through which miR-10b governs adipogenesis. Among the genes suppressed by miR-10b-5p, Gata6, an endoderm related gene (62), was predicted to bind miR-10b in the 3’ UTR (Figure 6A). We initially assessed the mRNA expression levels of Gata6 and its downstream target Bmp2 during adipogenesis (62). We observed that the expression pattern of both these genes was upregulated in response to miR-10b deletion (Figure 6B). To investigate if miR-10b directly targets Gata6, we treated miR-10b KO cells with a miR-10b mimic or a NTC miRNA mimic. Analysis by qRT-PCR showed that overexpression of miR-10b downregulated the expression of Gata6 (Figure 6D).

**Figure 6.**
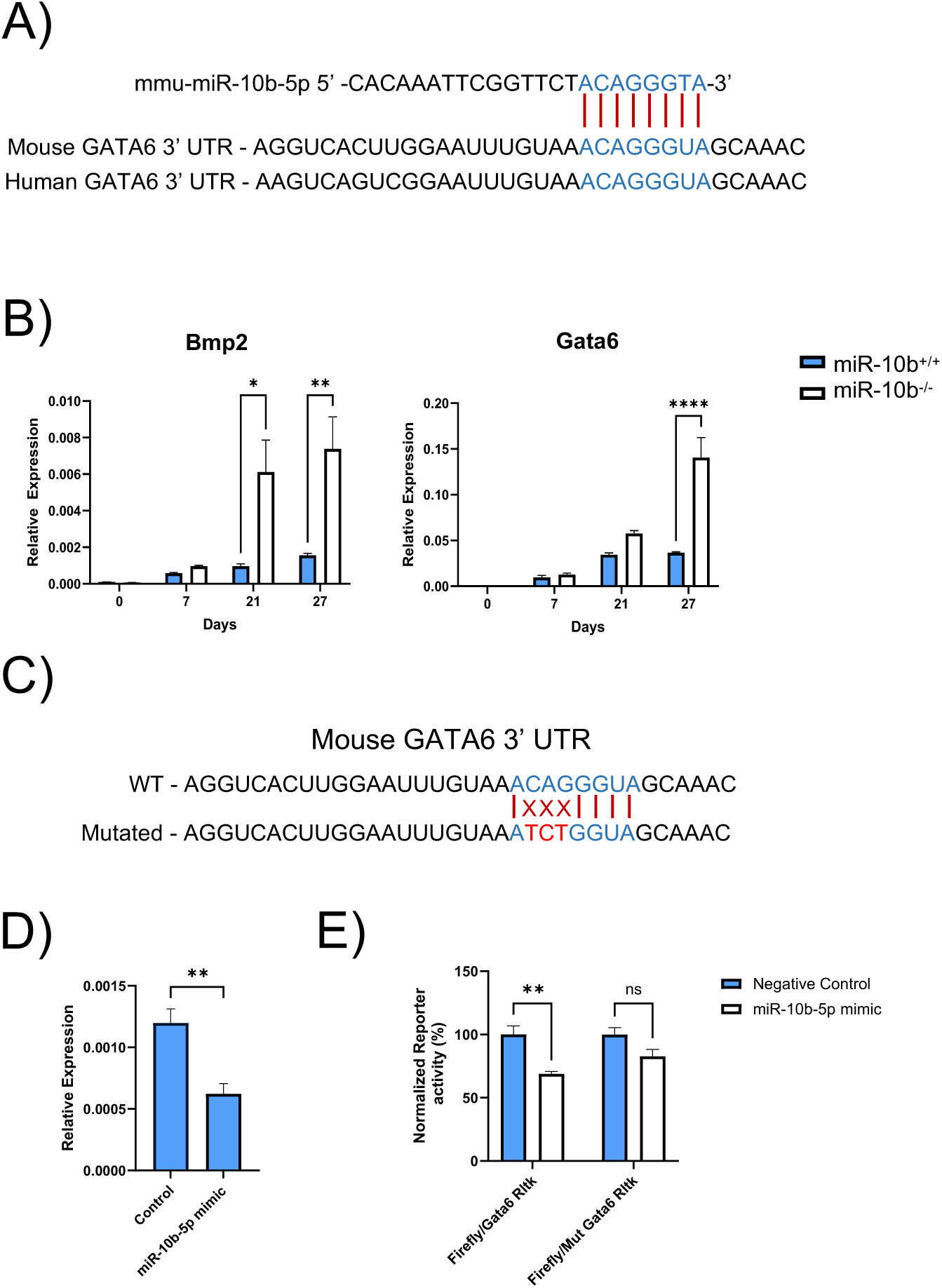
GATA6 as a potential target for miR-10b-5p. A) Binding seed sequence of miR-10b with the 3’UTR of Gata6. B) The relative expression of Bmp2 and Gata6 mRNA during mESC differentiation to adipocytes. C) The mutated seed sequences of Gata6-UTR was denoted in red. D) The expression of Gata6 was analysed by qRT-PCR in KO cells in response to miR-10b mimic. E) Luciferase reporter assays demonstrating the target relationship between miR-10b-5p and Gata6. The wild-type (WT) or mutant-type (MUT) constructs were inserted into the pRL-TK vector (Promega, UK) and firefly and Renilla luciferase activities were determined (n=3, mean ± SEM. ANOVA, **P<0.01, ns: not significant).

We generated luciferase reporters that had either a wild-type (WT) Gata6 3’UTR or a 3’UTR-containing mutated (MUT) miR-10b binding site (Figure 6C). Dual luciferase reporter analysis in HEK293 cells demonstrated that co-transfection of miR-10b-5p significantly suppressed reporter activity of the WT Gata6 3’UTR, but not with the mutated binding site (Figure 6E). Collectively, these results evidence that Gata6 is a direct target of miR-10b-5p.

### 6.8 Ablation of miR-10b-5p in Brown Adipocytes severely Impairs Differentiation

To further delineate the potential functions of miR-10b-5p during adipogenesis, we investigated if loss of function of this miRNA also influenced the later stages of differentiation. To this end, immortalized brown pre-adipocytes were treated with miRCURY miRNA Inhibitor that silenced miR-10b-5p and subjected cells to standard brown adipogenic treatment. Cells were also treated with a negative control that did not exhibit homology to any mouse sequence in the NCBI and miRBase databases. The silencing of miR-10b-5p in brown preadipocytes was initially confirmed through qPCR, demonstrating a reduction of its levels by over 100-fold when compared to control. This decrease persisted throughout the differentiation process until day 7 (Figure 7A).

**Figure 7.**
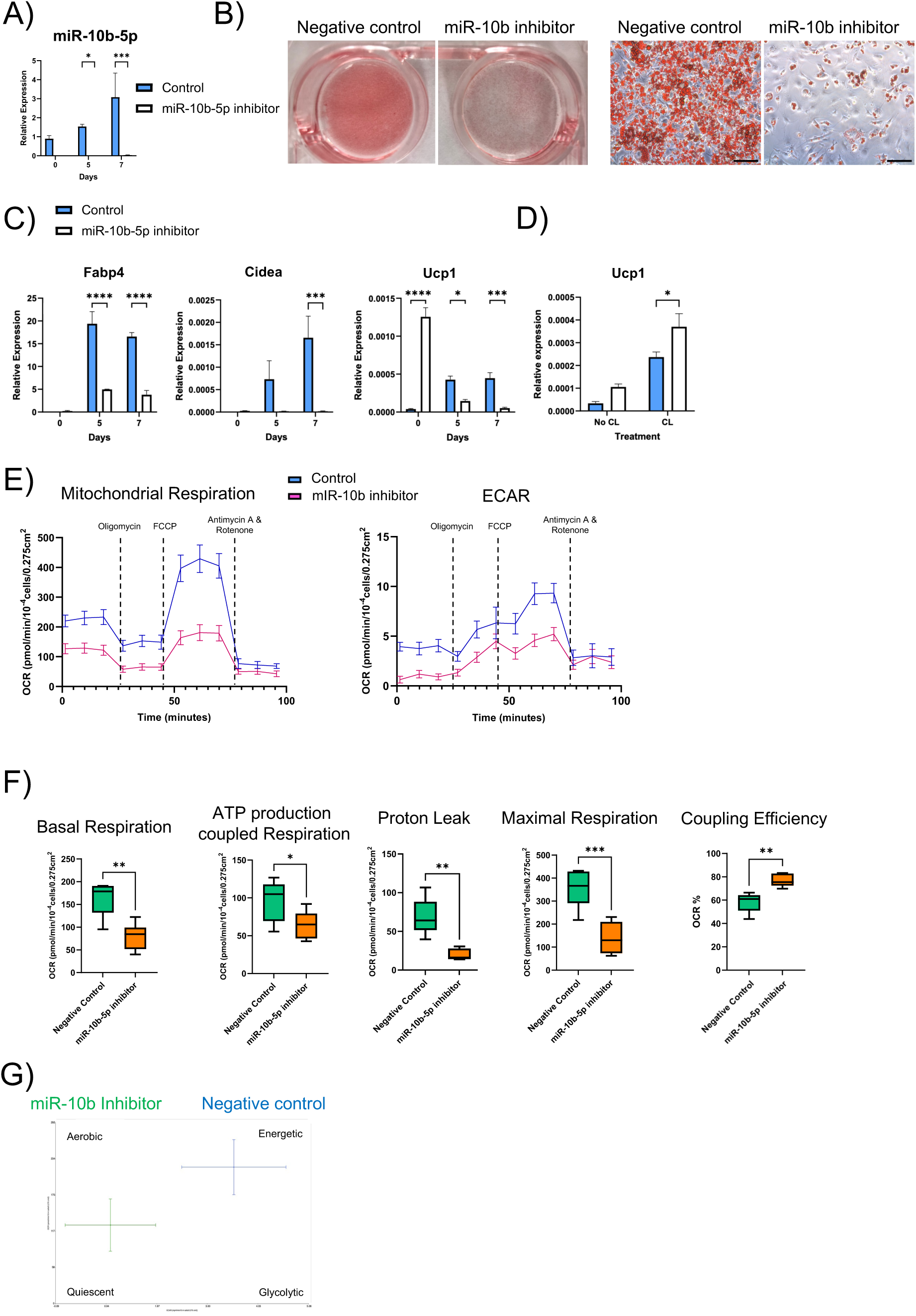
Downregulation of miR-10b severely compromised brown adipogenesis. A) miR-10b-5p levels were reduced in response to miR-10b-5p inhibitor. B) Representative Oil-Red-O staining of differentiated brown adipocyte cultures treated with miR-10b inhibitor or negative control. Scale bar: 50 μm. C) qRT–PCR analysis of gene expression of adipogenic and brown fat-selective genes in brown adipocytes. D) On day 8, mature BAs were treated with a miR-10b inhibitor or control for 48 h, followed by 4 h treatment with CL 316,243 or DMSO, and Ucp1 and Ppargc1a expression was analysed by qRT–PCR. Data were analysed with Student’s t-test and are presented as mean ± SEM (n ≥ 3). E) Effects on Bioenergetics triggered by miR-10b inhibition. Extracellular acidification rate (ECAR) and oxygen consumption rate (OCR) were measured eight days after transfection (n ≥ 5). Blue line: negative control, pink line: miR-10b-5p inhibitor. F) Box and whisker plots showing the components of mitochondrial respiration changes in differentiated BA, ATP production coupled respiration, Proton leak and maximal respiration capacity. Coupling efficiency was calculated by the ratio of ATP-coupled respiration and basal respiration, calculated as a percentage (%). G) Summary metabolic profiles as determined by plotting ECAR against OCR. Data were analysed with Student’s t-test and are presented as mean ± SEM (n ≥ 5). Centrelines indicate the medians and whiskers extend to the minimum and maximum values. (*p< 0.05, **p< 0.005, ***p< 0.0005, ****p < 0.0001).

Knockdown of miR-10b-5p in brown preadipocytes significantly suppressed brown adipogenesis as judged by reduction of Oil Red O Staining compared to cells treated with negative control (Figure 7B). This was also evidenced by the decreased expression of adipogenic and brown fat-selective genes (Figure 7C). Interestingly, Ucp1 expression was transiently upregulated prior to the treatment of standard adipogenic induction at day 0, reproducing the observation in miR10b-5p knockout ESCs. To specifically explore the functional effect of miR-10b in mature adipocytes, fully differentiated BAs were treated with miR-10b inhibitor or control for 48 hours. Activation of the beta 3 adrenergic receptor (β3AR) with the agonist CL 316,243 (CL) resulted in greater Ucp1 expression in cells treated with the miR-10b inhibitor compared with control cells (Figure 7D).

To examine the consequence of downregulation of miR-10b on bioenergetics, OCR and extracellular acidification rate (ECAR) in differentiated adipocytes at day 8 was measured by Seahorse assay (Figure 7E, F, G). miR-10b loss of function decreased basal respiration, respiration used for ATP production, mitochondrial respiration, proton leak and maximal respiratory capacity. Basal ECAR was also reduced by inhibiting miR-10b both before and after oligomycin treatment, and when analysed against OCR. The inhibition of miR-10b reduced glycolysis and oxidative phosphorylation, which is representative of fewer metabolically active cells, and increased coupling efficiency. We also investigated the functional impact of miR-10b-5p inhibition on bioenergetics in fully differentiated BAs. To achieve this, mature BAs were transfected with a miR-10b-5p or negative control inhibitor, and a Seahorse mito stress assay was performed 48 hours later. Knocking down miR-10b in mature adipocytes led to a reduction in maximal respiration, as measured by the OCR assay (Supplementary Figure S3). Consistent with the previous findings, basal respiration, ATP-linked respiration, mitochondrial respiration, proton leak, and maximal respiratory capacity were all decreased. Furthermore, coupling efficiency was increased in miR-10b-5p-inhibited cells. Collectively, these data suggest that reducing miR-10b-5p levels alters the mitochondrial metabolic profile.

### 6.9 Dynamic impact of miR-10b inhibition on different molecular processes that govern different stages of brown adipogenic differentiation

To unravel the pathways that are affected by miR-10b depletion, a transcriptomic analysis was performed on days 0, 5, and 8 of BA differentiation in the absence and presence of miR-10b inhibitor. The PCA plot clearly indicated that the transcriptomic profiles of the six sample groups were distinctly different (Supplementary Figure S4). The primary factor influencing the observed variance was the differentiation status of the cells. Furthermore, the plot also illustrated that the largest changes occur between days 0 and 5 or 8 in the cells subjected to the negative control.

Transcriptomic analysis identified 1601 differentially expressed genes at key time points of brown adipogenesis in the two treatment groups based on log2 fold change of ≥2 and adjusted p-values ≤0.05 (Figure 8A).

**Figure 8.**
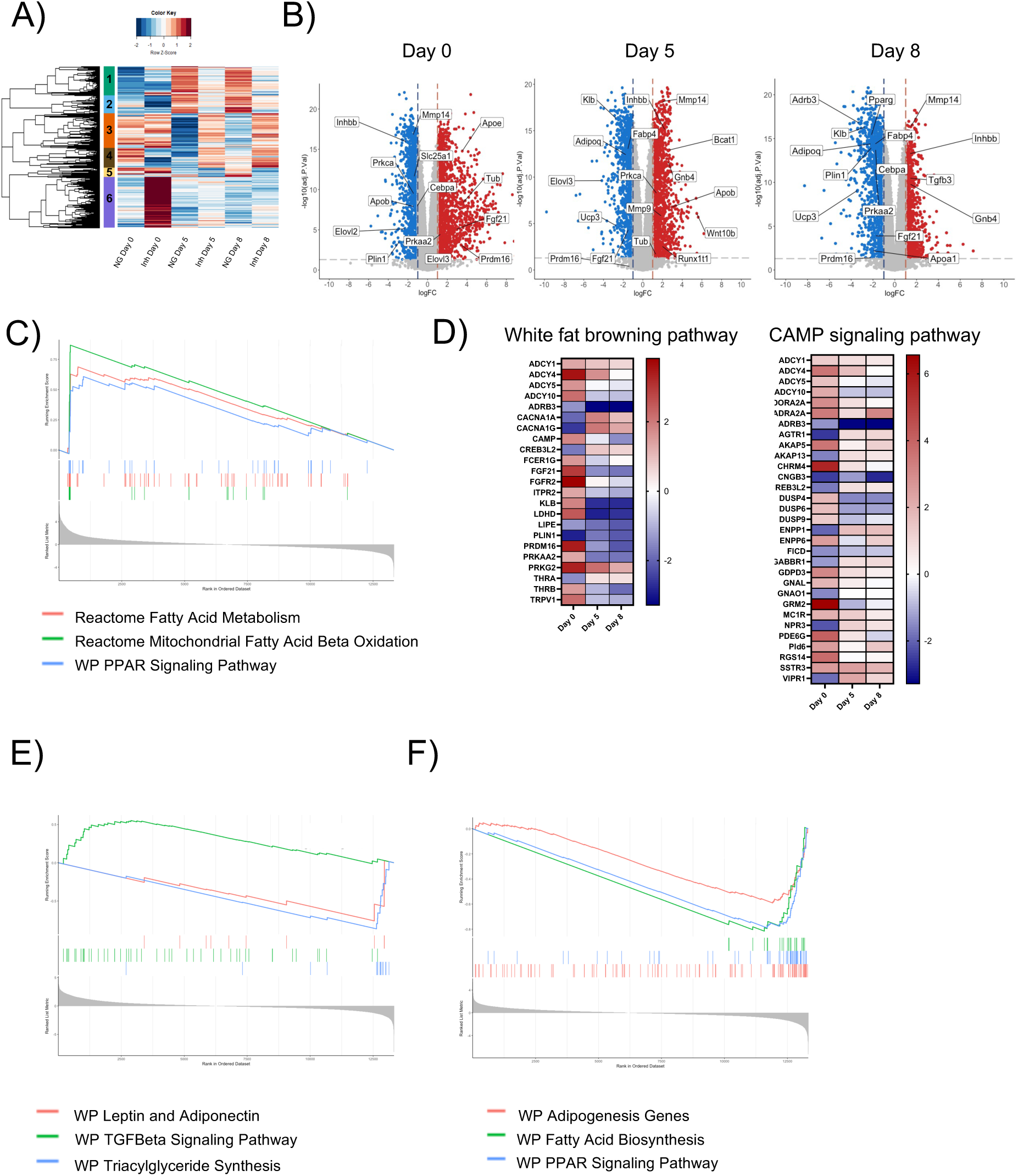
Transcriptomic analysis of brown adipocytes treated with miR-10b inhibitor or negative control during differentiation. A) Heatmap showing the Hierarchical cluster analysis of 1601 common DEGs between the treatment groups across the different time points of BA differentiation (log2(FC) > 2 and p adjusted value > 0.05). B) Volcano plots illustrating the expression profile of genes in BAs treated with miR-10b inhibitor (Inh) or NG at Day 0, 5 and 8 (log2(FC) > 1 and p adjusted value > 0.05). Negative logFC (red) = upregulated in BAs treated with mIR10b inhibitor; positive logFC (blue) = upregulated in BAs treated with NG. C) GSEA revealed a striking enrichment of transcripts involved in the Mitochondrial Fatty Acid Beta Oxidation, Fatty acid Metabolism and PPAR signalling among the genes that were more highly expressed during miR-10b inhibition at day 0. D) Heatmaps illustrating gene expression patterns associated with the White Fat Browning Pathway and cAMP Signalling Pathway identified through GSEA at day 0,5 and 8. E) GSEA demonstrated a reduction in the expression of genes related to Triglyceride Synthesis, as well as Leptin and Adiponectin, at day 5. F) GSEA results from day 8 dataset also exhibited a suppression in the levels of transcripts associated with Adipogenesis, PPAR Signalling Pathway and Fatty Acid Biosynthesis.

To compare the DEGs between the miR-10b inhibitor and control groups across the selected time points of differentiation, the adjusted p value ≤ 0.05 and the absolute value of log_2_ (fold change) ≥ 1 were employed as criteria. In total, 1730 genes were upregulated, and 1303 genes were downregulated at day 0, 1384 genes were upregulated, and 1143 genes were downregulated at day 5, whereas 956 genes were upregulated and 1318 downregulated in the late phase of BA differentiation (day 8) (Figure 8B). Specific genes associated with adipogenic differentiation (Apoe); thermogenic program (Fgf21, Prdm16) and lipid metabolism (Elovl3) were increased at day 0. In contrast, there were also genes related to the same pathways that were downregulated at day 0, including Cebpa, Prkca and Inhbb (Figure 8B). Additionally, a Gene Ontology (GO) analysis was conducted on cluster module 6 of the heatmap (Figure 8A), identified as the most densely populated module with the highest upregulated genes in cells treated with the miR-10b inhibitor compared to the negative control. Enrichment analysis showed that several enriched GO terms were related to cell development and differentiation, as well as genes associated with lipid metabolism. Additionally, GO analysis revealed enriched transcription factors, including Pparg, Smad3, and Srebf1 (Supplementary Figure S5).

GSEA of the day 0 dataset revealed a striking enrichment of transcripts involved in energy metabolism, lipid metabolism, and mitochondrial function, specifically within the BAs treated with miR-10b inhibitor (Supplementary Figure S6). The heightened expression of transcripts associated with Mitochondrial Fatty Acid Beta Oxidation, Fatty acid Metabolism and PPAR signalling suggests a role for miR-10b-dependent inhibition of metabolic processes critical for thermogenesis and fat metabolism in the undifferentiated state (Figure 8C, Supplementary Figure S6). We also observed a substantial enrichment of genes associated with the white fat browning pathway and the cAMP signalling pathway in BAs with miR-10b inhibition at day 0 compared to the other days (Figure 8D). These results demonstrate that key genes related to the brown adipogenic program are activated upon miR-10b inhibition at day 0. However, upon induction of the adipogenic differentiation program and during the late phase of differentiation, the majority of those key genes display markedly reduced expression with miR-10b inhibition. Several of these biological pathways overlap between the two datasets at day 5 and day 8 (Supplementary Figures S7 and S8). Upon mir-10b loss of function, biological pathways associated with lipid droplet formation, including those involved in triacylglycerol synthesis, fatty acid metabolism, and lipid metabolism, were suppressed during brown adipogenic differentiation. Notably, key signalling pathways related to the function and development of BAs were also repressed, such as oxidative phosphorylation, PPAR Signalling Pathway, and leptin and adiponectin pathways (Figure 8E, F). Consistent with the finding of perturbed differentiation, GSEA also revealed activation of the TGF-beta signalling pathway (Figure 8E), which previous research indicates hinders preadipocyte maturation (63,64).

Gata6 levels were upregulated during mouse and human BA differentiation (Supplementary Figure S12A, B). However, TPM levels of Gata6 expression during mouse BA differentiation showed no significant variation between control and miR-10b KD cells (Supplementary Figure S12D), suggesting that miR-10b does not regulate Gata6 expression in brown adipocytes.

### 6.10 Tub abundance is regulated by miR-10b-5p

In silico prediction (65) identified multiple hits for miR-10b-5p targeting of Tub, a transcription regulator that transmits signals from G protein-coupled receptors (GPCRs) and G proteins (66). In our analysis of whole-transcriptome RNA sequencing during BA differentiation time course, we noted a significant increase in genes related to G Protein Signalling through Tubby (Figure 9A). In agreement with the transcriptomic analysis, we found that Tub mRNA (Figure 9B) and protein levels (Figure 9C and D) increased in response to transfection of miR-10b-5p inhibitor in BAs. Collectively, the expression pattern of Tub is negatively correlated with miR-10b-5p expression during BA differentiation. To characterize the expression of Tub, we quantified expression levels in different fat depots and cell types. Tub mRNA was significantly higher in classical BAT compared to gonadal and subcutaneous WAT in the mature adipocyte fraction (Figure 9E). Tub mRNA was consistently higher in the SVF compared to the adipocyte fraction, while maintaining the observed tissue-specific pattern (Supplementary Figure S12C). The effect of adipogenesis on Tub mRNA expression was assessed from BA cells at various key time points of differentiation. Tub was most abundant in confluent pre-adipocytes and at the onset of differentiation (day 0). However, its levels gradually declined as differentiation proceeded (Figure 9F). To gain a better insight into the cellular distribution of Tub protein during brown adipogenesis, its localization was visualized at different time points using confocal microscopy. On Day 0, Tub protein was primarily concentrated in the perinuclear region. By day 1, BAs displayed a uniform increase in Tub protein throughout the cell, with a fine punctate pattern distributed intracellularly. In the late stages of BA differentiation, Tub protein was only localised in the nucleus (Figure 9G). Together, these data reveal that Tub displays a distinct spatiotemporal dynamic pattern during brown adipogenesis.

**Figure 9.**
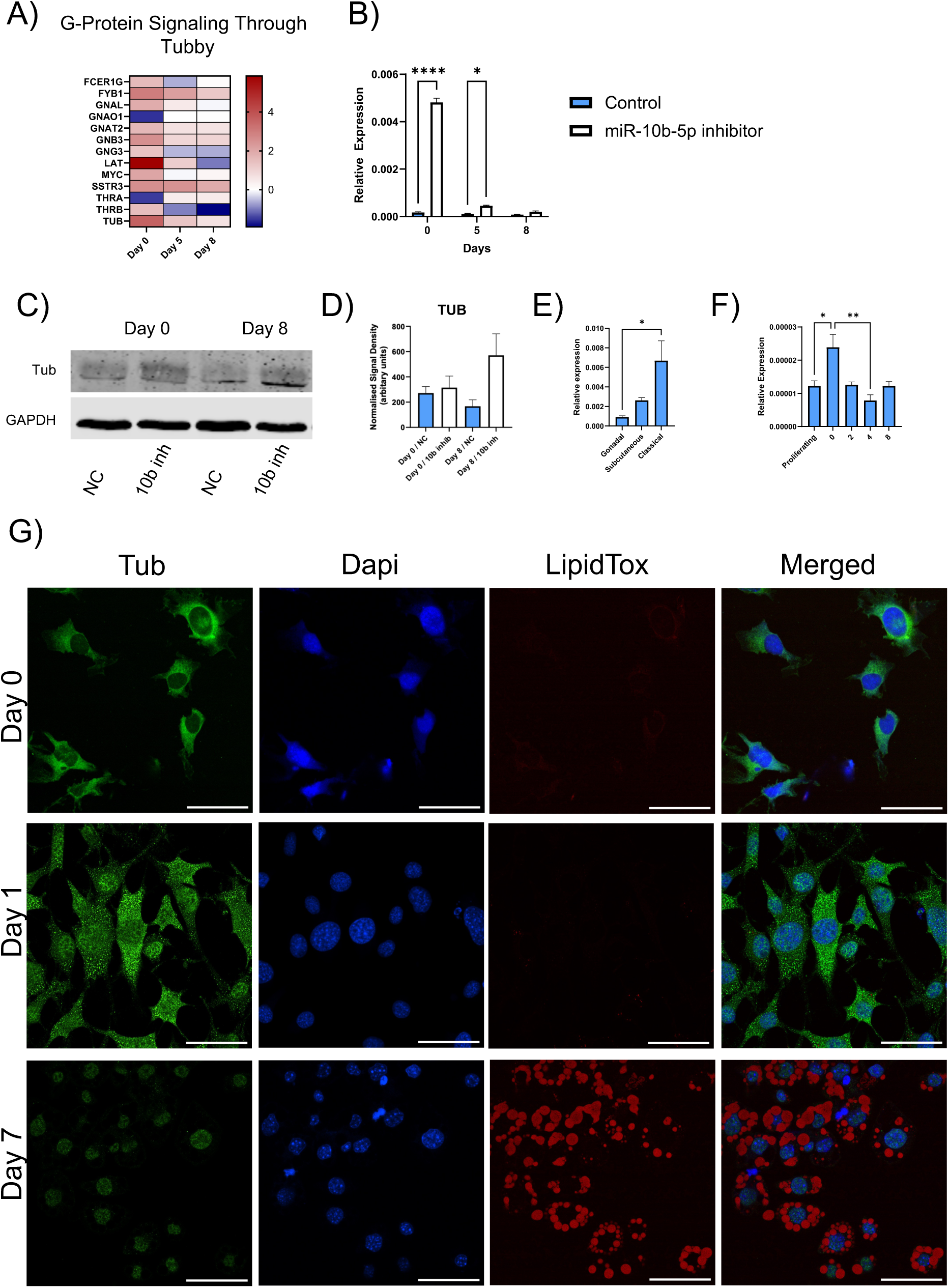
The expression level of Tub is negatively correlated with miR-10b-5p abundance during BA differentiation. Heatmap depicting gene expression patterns linked to G-protein Signalling through Tubby identified through GSEA at day 0,5 and 8. B) qRT– PCR analysis of Tub expression in response to miR-10 b-5p inhibitor treatment at specific time points of BA differentiation. C) Protein levels of TUB were analysed at pre-adipocyte (day 0) and fully mature adipocyte state (day 7) in response to miR-10b-5p or negative control treatment. D) Quantification of TUB levels normalized by GAPDH, in units relative to peak intensity. E) Measurement of TUB expression in three different adipose tissue depots: adipocyte-fraction from gonadal WAT, subcutaneous WAT and classical BAT. mRNA levels were standardised against those for L19. (2-way Anova, n=3, *p<0.05). F) RNA was harvested from BA cells at different stages of differentiation and Tub mRNA levels quantified by RT-qPCR. (2-way Anova, n≥4, *p<0.05, **p<0.005). G) Tub immunostaining during BAT differentiation. Scale bar: 50 µm. Images taken at 40x magnification. NC: Negative control, 10b inh: miR-10b-5p inhibitor.

### 6.11 Tub is a direct target of miR-10b-5p

To investigate whether miR-10 b-5p can directly target Tub mRNA we cloned a section of the 3′UTR segment of Tub containing the predicted miR-10 b-5p target site or with the seed site mutated (Figure 10A) into the RLTK luciferase reporter construct. The reporter constructs were co-transfected into HEK293 cells with either miR-10b-5p or negative control mimics. Luciferase reporter assays showed that, unlike the negative control, miR-10 b-5p mimics reduced the activity of the reporter construct harbouring a wild-type 3′UTR (Figure 10B). In contrast, mutations in the seed sequence prevented miR-10 b-5p-dependent suppression of reporter gene activity. This demonstrates that miR-10b-5p directly interacts with the predicted target sites in the Tub transcript.

**Figure 10.**
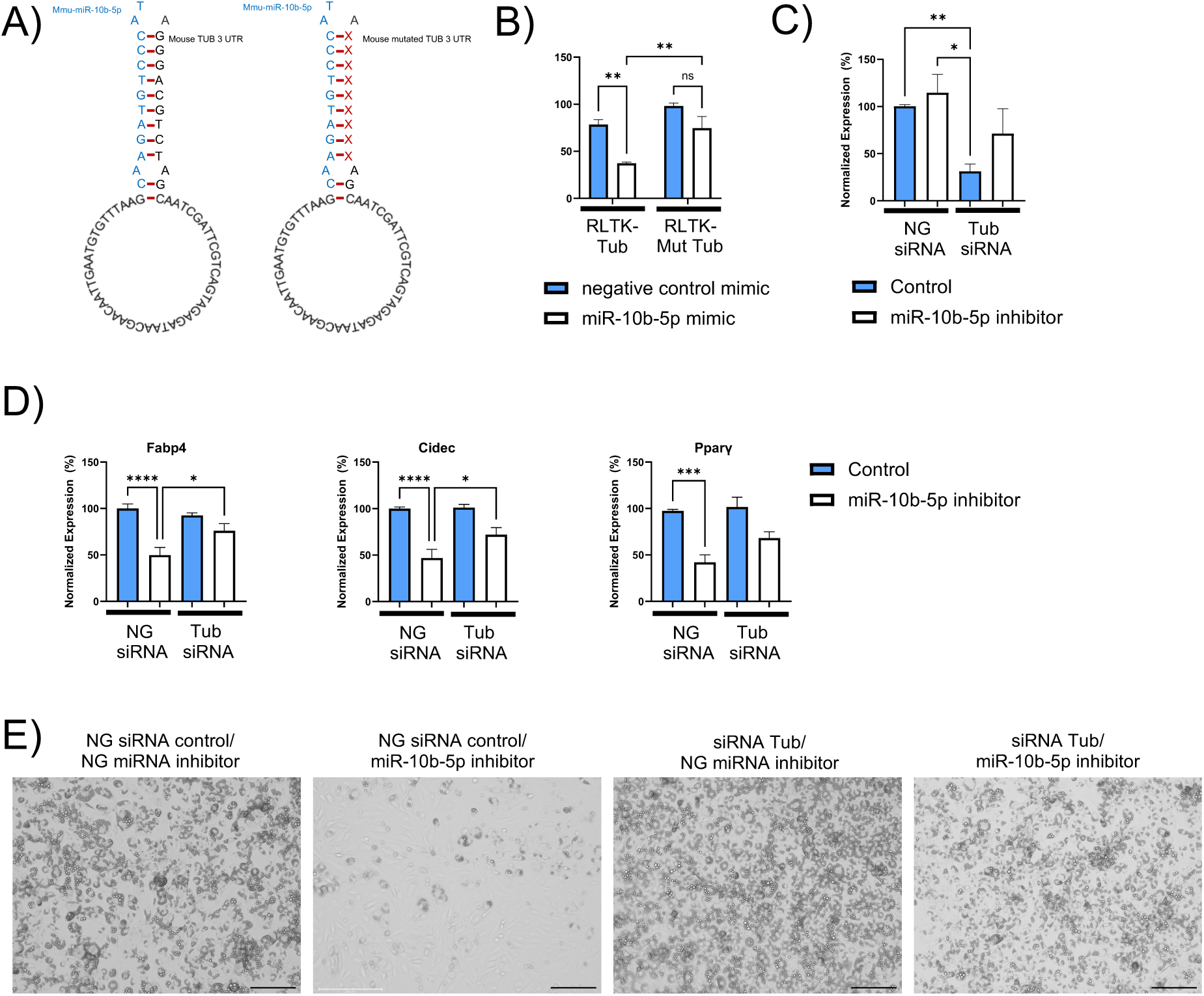
Tub is a direct target of miR-10b-5p. A) Binding seed sequence of miR-10b with the 3’UTR of Tub. The mutated seed sequences of Tub-UTR are denoted in red. B) Luciferase reporter assay demonstrating the target relationship between miR-10b-5p and Tub (n≥3, mean ± SEM. 2-way ANOVA, *P<0.05, **P<0.005). C) Tub expression levels in mature BAs were measured after transfection with either negative control siRNA and miR-10b-5p inhibitor/miRNA negative control inhibitor, or Tub siRNA and miR-10b-5p inhibitor/miRNA negative control inhibitor prior to adipocyte differentiation (Kruskal-Wallis test, n≥8, *p<0.05, ***p<0.0005). D) RT-qPCR analysis reveals the gene expression signatures of Fabp4, Cidec, and Pparγ in differentiated BAs, which were initially treated with either a miR-10b inhibitor or negative control and co-transfected with siRNA targeting Tub or a siRNA negative control prior to differentiation. (Fabp4 and Cidec were analysed with 2-way Anova, n≥8, *p<0.05, ****p<0.0001, PPARγ were analysed with Kruskal-Wallis test, n≥12, ***p<0.0005). E) Representative bright-field images show the level of BA differentiation by visualizing lipid droplet accumulation on day 6. Magnification 10x. Scale bar 100 µm. NG: Negative Control

To further investigate this finding, BAs were transfected with miR-10b-5p inhibitor in the presence or absence of Tub siRNA and then subjected to the standard differentiation protocol. Tub mRNA was similar in fully differentiated cells exposed to miR-10b inhibitor or its control prior to differentiation (Figure 10C), in agreement with the earlier time course (Figure 9B). Tub siRNA decreased Tub mRNA expression, although levels were higher in cells co-transfected with miR-10b-5p inhibitor. This indicates that inhibiting miR-10b can counter the action of siRNA on Tub expression at the later stages of differentiation.

To investigate the suppression of BA differentiation, we monitored expression of key genes associated with adipogenesis and examined morphological changes in cells exposed to combinations of miR-10 b-5p inhibitor and Tub siRNA. Knockdown of Tub partially reversed the miR-10 b-mediated decrease in Fabp4, Cidec and Pparg upon differentiation (Figure 10D), indicating that the increase in Tub expression due to miR-10 b-5p inhibition appears to be essential for the observed effects. This was further evidenced by the decreased lipid droplet formation in cells treated with the miR-10b-5p inhibitor on day 6, being mitigated by co-transfection with Tub siRNA Tub (Figure 10E, Supplementary Figure S9).

### 6.12 Increased abundance of miR-10b-5p in white adipocytes enhances adipogenesis

As miR-10b-5p was found to be significantly enriched in BAs in comparison to WAs, we investigated the effects of miR-10b-5p upregulation on WA differentiation. Immortalized white pre-adipocytes were transfected with hsa-miR-10b-5p miRCURY LNA miRNA Mimic or Negative Control mimic and subjected to standard white adipogenic differentiation. MiR-10b-5p mimic significantly enhanced white adipogenesis as judged by the increased production of lipid droplets in cells at day 3 and 6 compared to negative control (Figure 11A). Interestingly, by day 8, the negative control group appeared to have reached similar levels of lipid droplet synthesis as the mimic group, as shown by staining for lipid droplets (Supplementary Figure S10). However, the mimic group appears to have larger lipid droplets compared to the negative control (Figure 11A). The elevated expression of adipogenic marker Fabp4 further strengthened this observation. We also observed increased Ucp1, Cidea and Ppargc1a levels during white adipogenesis, indicating a possible increase in the expression of a thermogenic gene program (Figure 11B). While increased levels of miR-10b-5p in WAs elevated their rate to undergo differentiation, we tested whether the cells also gained propensity for β-adrenergic stimulation. Activation of the β3AR with CL resulted in a robust increase of Ucp1 in cells, regardless of the treatment (miR-10b mimic or control). (Figure 11C, D). We also investigated Tub expression during WA differentiation in response to mimic treatment. Downregulation of Tub due to increased levels of miR-10b-5p was observed at the mRNA level on day 0 (Figure 11E), whereas downregulation at the protein level was seen on day 3 (Figure 11F, G). It is noteworthy that the Tub protein demonstrated a marked transient increase in expression on day 3 of differentiation. Tub mRNA was undetectable at days 5 and 8 (data not shown). To determine whether functional changes in bioenergetics accompanied the observed thermogenic-induced effects of miR-10b-5p mimic, differentiated WA cultures were analysed using the Seahorse assay (Figure 11H-J). The miR-10b-5p mimic during WA differentiation instigated a substantial increase in maximal respiration, signifying an enhanced capacity for oxygen consumption under conditions of elevated energy demand. While not statistically significant, the treatment appeared to enhance ATP production–coupled respiration, suggesting a potential improvement in the efficiency of ATP synthesis. Moreover, there was also a rise in proton leak, indicating a higher mitochondrial uncoupling, which could potentially contribute to thermogenesis. Remarkably, coupling efficiency, which means the ratio of ATP-coupled respiration and basal respiration, also exhibited a rise, suggesting a more efficient coupling of mitochondrial respiration to ATP synthesis. Collectively, these results highlight that the miR-10b-5p mimic improved various parameters of mitochondrial bioenergetics, potentially contributing to increased cellular energy metabolism and function.

**Figure 11.**
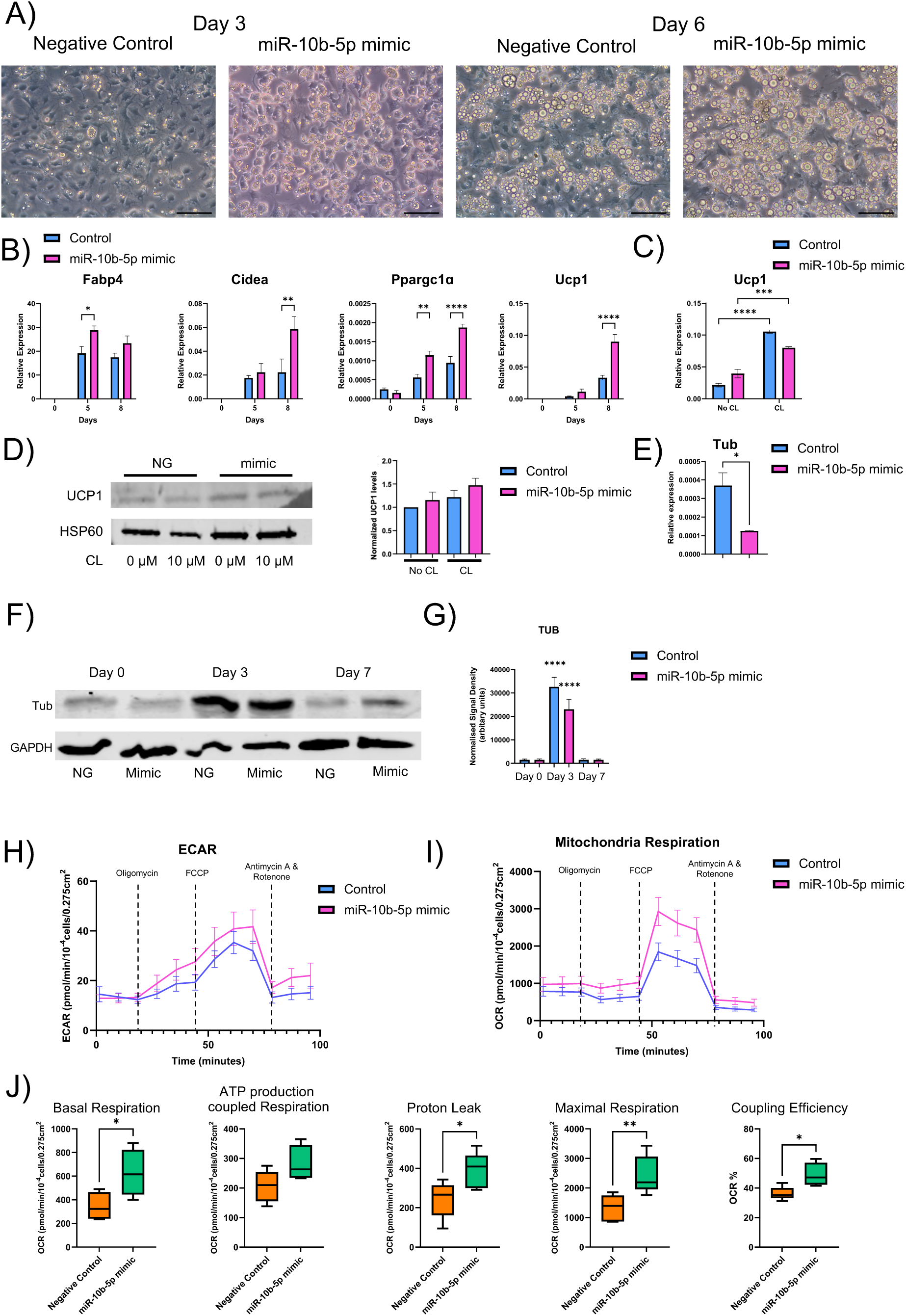
Higher levels of miR-10b-5p in white adipocytes promote adipogenesis. A) The rate of differentiation was assessed by observing the level of lipid accumulation on days 3 and 6. A representative bright-field image was captured from cultures treated with either negative control or mimic, depicting the progress of differentiation. B) C) qRT–PCR was performed to analyse the gene expression of adipogenic and brown fat-selective genes in white adipocytes (2-way Anova, n≥3, *p<0.05, **p<0.005, ***p<0.0005, ****p<0.0001). C) On day 8, cells were treated with 10 μM CL 316,243 (CL) or DMSO (control) for 24 hours. UCP1 mRNA was then analyzed by qRT-PCR (2-way Anova, n≥3, ***p<0.0005, ****p<0.0001). D) Cell lysate was also isolated for protein analysis by western blotting. E) Tub mRNA was decreased in response to miR-10b-5p mimic treatment at day 0 (t-test, n≥3, *p<0.05). F) Immunoblots displaying protein levels of Tub during key time points of WA differentiation. G) TUB protein levels were quantified and normalized against housekeeping protein GAPDH (2-way Anova, n≥3*p<0.05, **p<0.005, ***p<0.0005, ****p<0.0001). Data are presented as mean ± SEM. H) Effects on Bioenergetics triggered by overexpression of miR-10b-5p. ECAR and I) OCR were measured at day 8 (n ≥ 4). J) Box and whisker plots depicting the components of mitochondrial respiration changes in differentiated WA, ATP production coupled respiration, Proton leak and maximal respiration capacity. Coupling efficiency was measured by the ratio of ATP-coupled respiration and basal respiration, calculated as a percentage (%). Centrelines indicate the medians and whiskers extend to the minimum and maximum values. (Student’s t-test, n ≥ 5, *p < 0.05, **p < 0.01).

To gain more insight into Tub’s mechanisms during the process of white adipocyte differentiation, we stained white adipocytes at various stages of differentiation (Figure 12). Tub expression was elevated during the early stages of differentiation, contrasting with reduced expression in later stages.

**Figure 12.**
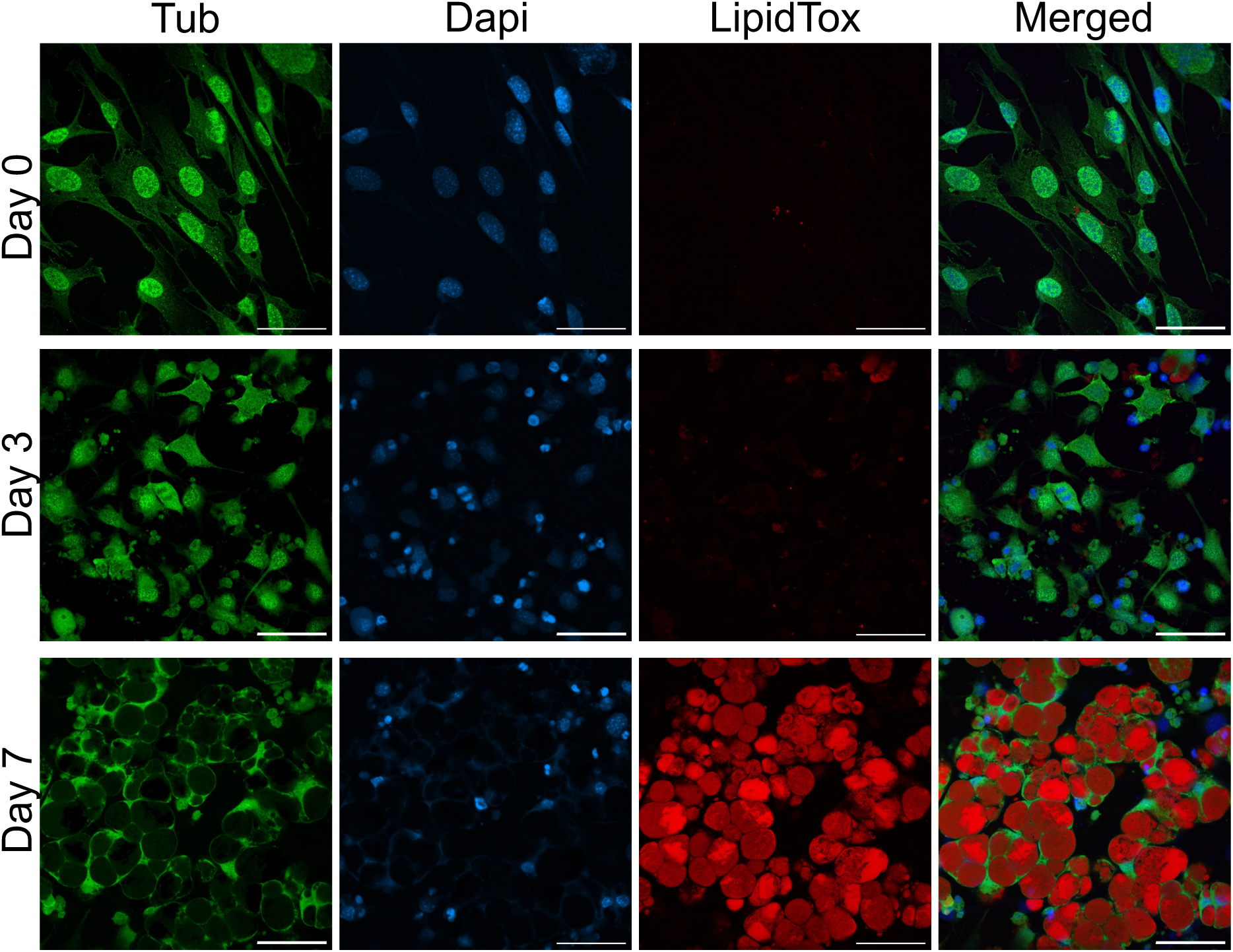
Tub immunostaining during WAT differentiation. Scale bar: 50 µm.

Notably, on day 0, Tub exhibited strong nuclear localization, whereas on day 7, it was more uniformly distributed throughout the cytoplasm. Additionally, Tub protein displayed a punctate pattern on day 0, although to a lesser extent compared to BAs. On day 3, tub protein appears to be localised in the perinuclear region. We also studied Tub protein behaviour in response to miR-10b-5p mimic. The negative control exhibited uniform Tub staining throughout the cell, whereas miR-10b-5p overexpression led to a punctate pattern, hinting at a distinct cellular response. Fluorescence intensity remained consistent across treatments, yet variations in lipid droplet sizes were evident (Supplementary Figure S11). These findings emphasise Tub’s dynamic involvement in adipocyte differentiation, indicating intricate regulatory processes that warrant further investigation.

## 7 Discussion

Adipogenesis is a highly complex process and is dynamic in nature. It involves the transition of pre-adipocytes into nondividing adipocytes and the key mechanisms that promote preadipocytes commitment and maturation includes multiple epigenetic factors, miRNAs as well as protein regulators. However, it is still unknown how and when an irreversible conversion occurs into a definite state capable of accumulating lipids (67). With the ongoing obesity epidemic and the strong associations of obesity with diabetes, cardiovascular disease and carcinogenesis (68,69), understanding the molecular processes underlying adipogenesis, or the transition of dividing preadipocytes into nondividing, lipid-accumulating fat cells, is of pivotal importance and therapeutic interest. Substantial evidence has demonstrated that miRNAs play a crucial role in mouse early embryonic development and may direct germ layer specification (70). In the current study, we found that miR-10b-5p is a positive regulator in adipocyte lineage commitment and early adipogenesis. Our findings strongly suggest that its depletion hinders the commitment steps of the adipogenesis process, which involves the conversion of ESCs into preadipocytes. Transcript analysis with qRT-PCR and RNA sequencing conducted at specific time points during adipogenesis in both the KO and NTC groups demonstrated elevated expression of pluripotency markers (Nanog, Oct4 and Sox2) and endoderm markers (Sox7, Gata4) in the KO group. In agreement, GSEA revealed a significant enrichment of transcripts linked to pluripotent mechanisms, alongside a potential readiness towards committing to the endodermal lineage. Furthermore, genes involved in BMP signalling, such as Smad9 and Bmp5 (59,60), were also downregulated in the KO group, which could lead to a reduction of BMP signalling, which is crucial for mesodermal commitment (61). The GATA family of transcription factors are considered critical for embryonic development, playing fundamental complex roles in cell fate decisions and tissue morphogenesis. Among the GATA factors, Gata4 and Gata6 are recognized for their significance in cell types derived from the endoderm (71). GATA6 is essential for primitive endoderm specification, as embryos lacking Gata6 default all inner cell mass cells to epiblast fate and fail to survive implantation. Furthermore, Gata6 heterozygous embryos have delayed primitive endoderm development due to reduced Gata6 levels (72). Previous research also demonstrated that Nanog and Gata6 co-bind to the same regulatory elements, and their relative levels play a critical role in determining cell fate during early embryonic development (73). In the present study, we found that Gata6 is a direct target of miR-10b-5p through binding-site prediction and dual-luciferase assay validation. Gata6 was down-regulated by treating KO cells with miR-10b-5p mimic and had an inverse correlation with miR-10b-5p mimic and had an inverse correlation with miR-10b-5p expression during adipogenesis. Previous research has shown that ectopic overexpression of Gata6 in mESCs induced the development of visceral endoderm, while also upregulating the expression of Bmp2. Conversely, deletion of Gata6 prevents embryonic stem cells from differentiating into endoderm (74). This highlights the critical regulatory role of Gata6 in directing cell fate decisions during early embryonic development (75). Its expression during brown fat development has not been documented thus far (76). Nevertheless, *in silico* analysis identified Gata6 as a potential transcription factor that may modulate the expression of brown fat-specific genes such as Zic1 and enzymes involved in the tricarboxylic acid (TCA) cycle, including citrate synthase (Cs), as documented in the ProFat database (77). In this study, miR-10b-5p potentially directs cells towards the mesoderm lineage and facilitates their commitment to pre-adipocytes by controlling the expression of Gata6 and its downstream target Bmp2. Considered together, these findings offer the first line of evidence that miR-10b-5p might modulate stem cell fate decision, thereby establishing a foundation for deeper investigation into the molecular mechanisms underlying miRNA influence on lineage commitment (Figure 13). Future studies are warranted to fully define the role of miR-10b-5p as a stem cell fate switch and provide fundamental insights into early mammalian cell fate specification.

**Figure 13.**
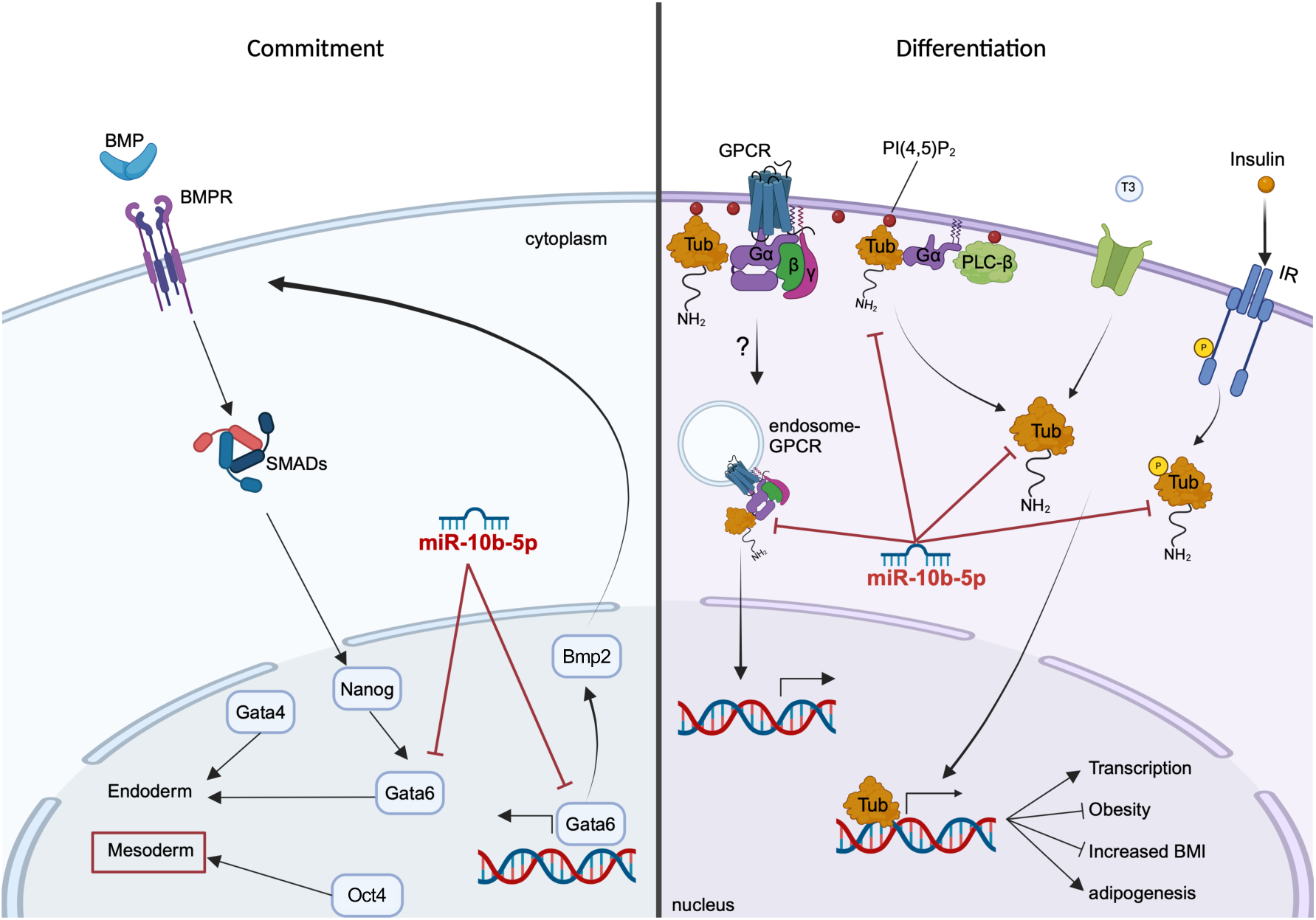
A Model of miR-10 b-5p-mediated signalling driving lineage commitment and adipogenesis. During stem cell differentiation, BMPs initiate signalling through binding to BMP receptors (BMPRs), leading to phosphorylation and activation of Smad1, Smad5, and Smad8. These activated SMADs translocate into the nucleus to regulate gene expression that promotes mesoderm and endoderm differentiation. In mESCs, Nanog binds Smad1 and blocks coactivator recruitment, limiting BMP signalling and mesodermal progression. Nanog also directly represses Gata6, which in turn downregulates Gata4, blocking transcriptional networks associated with primitive endoderm differentiation. Gata6 is required for cells to respond to lineage-inductive cues, which leads to downregulation of Nanog and progression toward a primitive endoderm fate. The balance between Nanog and Gata6 expression levels within individual cells serves as a key determinant in guiding the choice between embryonic and extraembryonic lineages. Additionally, Gata6 is a direct target of miR-10b-5p, and this post-transcriptional regulation may further influence the shift toward endodermal commitment rather than towards adipogenic lineage. Furthermore, miR-10b-5p was also identified as a positive regulator in the late stages of adipogenesis by directly regulating Tub expression. Tub expression is regulated by T3, and it functions as a membrane-bound transcription regulator that translocates to the nucleus upon phosphoinositide hydrolysis, thereby linking GPCR signalling to gene expression. Insulin induces its tyrosine phosphorylation and promotes its association with the insulin receptor.

In the present study, miR-10b-5p was also identified as a positive regulator in the late stages of adipogenesis, evidenced by a reduction of lipid droplet accumulation and decreased adipogenic gene expression in the presence of miR-10b inhibitor. This halt in adipocyte differentiation into mature adipocytes was accompanied by alterations in the mitochondrial metabolic profile and the presence of reduced metabolically active cells. Interestingly, miR-10b-5p did not appear to influence Gata6 expression in brown adipocytes. However, RNA sequencing identified a significant upregulation of genes linked to G protein signalling associated with increased Tub expression. In line with our transcriptomic results, Tub mRNA and protein levels increased with miR-10b-5p inhibition in BA and declined with miR-10b-5p overexpression in WA. Reporter gene assay indicated that miR-10b-5p directly regulated Tub expression. Moreover, silencing Tub partially reversed the inhibitory effect of miR-10b-5p in the differentiation of brown adipocytes. Mutations in the Tub gene are known to lead to late-onset obesity and insulin resistance in mice (78–81) as well as syndromic obesity in humans (82). Gene knockout studies of Tub in mice or *C. elegans* produce a similar phenotype as the Tubby mouse seen at the Jackson Laboratories, indicating that the phenotype is due to a loss-of-function mutation (80,83). Furthermore, adipogenesis plays a critical role in regulating healthy adipose tissue distribution and remodelling in obesity (84). If adipogenesis is impaired during excessive energy intake, then existing adipocytes continue to undergo hypertrophy to store excessive energy, leading to metabolic dysfunction (85). Taking into account our observations on the miR-10b-5p and Tub expression during adipocyte differentiation and previously reported association between the Tub and increased risk of obesity (86,87), we hypothesise that the required transient increase in Tub expression during the early stages of adipogenesis must be regulated by miR-10b-5p expression for adipogenesis to proceed appropriately (Figure 13).

Despite Tub’s clear physiological relevance to obesity, its precise molecular function remains unclear. Protein structure analysis of Tub revealed a potential DNA-binding groove in its COOH-terminal DNA-binding domain, along with possible transcriptional activation domains in its N-terminal region, strongly indicating that TUB functions as a transcription factor (87). Santagata et al. (88) demonstrated that TUB protein likely acts as a membrane-bound transcription regulator that translocates to the nucleus in response to phosphoinositide hydrolysis, establishing a direct link between heterotrimeric GTP-binding protein (G protein)-coupled receptors and gene expression regulation. Specifically, Tub localizes to the plasma membrane by binding phosphatidylinositol 4,5-bisphosphate through its carboxyl terminal. Receptor-mediated activation of G protein α_q_ (Gα_q_) induces PLC-β activity, causing TUB to detach from the plasma membrane and translocate to the nucleus (88). Notably, transfecting Neuro-2A or HEK293 cells with GFP-Tubby fusion proteins revealed that, after initially localizing to the plasma membrane, fluorescence began accumulating in the nucleus after 36 hours. This nuclear accumulation was primarily observed in cells with high expression levels. These findings suggest that TUB may translocate to the nucleus when its membrane binding sites become saturated (88).

Further studies in neuronal cells reinforce this hypothesis, showing that activation of the 5HT-2C serotonin receptor induces TUB nuclear translocation (88). In line with these findings, our immunostaining results demonstrated that Tub localisation differs throughout adipocyte differentiation as well as between different adipocyte depots. In brown preadipocytes, TUB is primarily localized in the cytoplasm. However, during adipogenic induction, as levels increase, its distribution becomes more uniform throughout the cell. In fully differentiated brown adipocytes, TUB localization shifts entirely to the nucleus. WA exhibited strong nuclear localisation, to a lesser extent, in the cytoplasm. Notably, the cytoplasmic presence of TUB persisted throughout WA differentiation. Two transcript variants encoding distinct Tub isoforms have been identified, both sharing the C-terminal domain but differing in their alternatively spliced NH2-terminal regions (80,88). The COOH-terminal domain of Tubby directs its localization to the plasma membrane, while the NH2-terminal domain directs it exclusively to the nucleus. The functional nuclear localization signal (NLS) is mapped to the sequence K_39_KKR within the NH2-terminal domain (88,89). Therefore, the observed variation in Tub localization might also result from differential splicing. It remains to be determined whether the different isoforms contribute to cell-specific expression of the TUB protein, which can be tested using epitope-specific antibodies. Interestingly, BAs exhibited a distinct fine punctate pattern intracellularly, while WAs showed this to a lesser extent. These structures may represent endosomes, supporting the emerging concept of alternative GPCR signalling via G proteins in intracellular compartments, such as early endosomes, rather than being confined to the plasma membrane (90). However, further research through endosome staining and co-localization with TUB is needed to confirm this. Furthermore, it would be of value to explore which GPCRs TUB interacts with and whether these interactions vary depending on different stimuli or stages of adipocyte differentiation. Remarkably, previous research has also implicated TUB as being integral to the trafficking of proteins to the primary cilium, a 1- to 10-μm-long antenna-like projection that extends from the plasma membrane. Primary cilia sense extracellular cues via receptors and relay signals from pathways like GPCRs, RTKs, Hedgehog, and Wnt, regulating an array of cellular processes including cell proliferation, migration, and differentiation (91–93). Recent studies have shown that primary cilium dysfunction influences adipogenesis (94,95). During adipocyte differentiation, the cilium appears in confluent cells, elongates early on, and disappears as lipid accumulation increases (96), mirroring the Tubby expression pattern we observed in brown adipocytes. Given that GPCR trafficking in and out of the cilium is tightly regulated by the Tubby protein family, these findings highlight a potential role for TUB in coordinating ciliary signalling during adipocyte differentiation.

Previous research in rodent adipocytes and neuronal cells has shown that Tub expression is regulated by insulin and T3, two key hormones involved in the regulation of metabolism (97,98). Insulin treatment of CHO-IR and PC12 cells induces tyrosine phosphorylation of Tub and facilitates its interaction with SH2 domain-containing proteins (99). Furthermore, cell sensitivity to insulin appears to regulate Tub expression in adipocytes (97). Stretton et al. (97) demonstrated that 3T3-L1 adipocytes exposed to chronic high insulin develop insulin resistance and show increased Tub levels, which are reduced by the insulin-sensitizing drug rosiglitazone. Consistently, we observed higher Tub expression in early differentiation, followed by a decline throughout the process, even when the miR-10b inhibitor was present. The observed reduction in Tub during adipogenesis is likely, at least in part, due to insulin-induced repression of the Tub gene. Insulin suppresses the expression of several genes encoding proteins with key metabolic functions (100) and Tub may be part of the group of repressed genes.

Notably, miR-10b-5p knockout ESCs, as well as BAs treated with the inhibitor, exhibited increased Ucp1 at day 0, resulting in elevated cellular respiration. This was also evident by the increased expression of transcripts associated with Mitochondrial Fatty Acid Beta Oxidation, Fatty acid Metabolism and PPAR signalling at day 0 in BAs (101). A transient increase in PPARG and UCP1 expression in human subcutaneous SVF cells was previously observed during differentiation. Similarly, another study found that UCP1 and PPARG peaked at day 14 and declined by day 28 post-differentiation in SGBS adipocytes (102), suggesting that cells may transiently adopt a brown adipocyte-like phenotype while differentiating to white adipocytes. Our results show that miR-10b depletion in cells may temporarily activate this process that would otherwise suppress UCP1 and Pparg, as observed in brown preadipocytes and fully differentiated BAs treated with the inhibitor.

We also examined whether increased miR-10b could induce a browning phenotype in white adipocytes. Mimic treatment enhanced adipogenesis and thermogenic effects as seen by increased Ucp1 and Ppargc1a, alongside bioenergetic changes. Collectively, our findings suggest a role for miR-10b in metabolic processes critical for lipid and energy metabolism. Pathogenesis of obesity has been linked to reduced levels of miR-10b-5p in mouse and rat (103). Furthermore, Herrera et al. (104) found decreased miR-10b expression in response to hyperglycemia, suggesting its role in type 2 diabetes pathophysiology in the Goto–Kakizaki rat model. This pattern was consistent with the miR-10b-5p expression profile observed in children with type 1 diabetes (T1D) (105), twins with type 2 diabetes (T2D) (106) and T2D patients (107,108), strengthening the critical role of miR-10b-5p during metabolic dysfunction. Singh et al. (109) further showed that miR-10b loss in β-cells and interstitial cells of Cajal (ICC) triggered diabetes and GI dysmotility in male mice via the miR-10b-KLF11-KIT pathway, which regulates glucose and gut motility. This effect was absent in female mice, highlighting hormonal differences in T2D susceptibility. They later found that in female T2D mice, characterized by insulin resistance, the pathogenesis was driven by the miR-10a/b-5p-NCOR2-INSR axis, and miR-10a/b-5p treatment improved glycaemic control. The *in vivo* relevance of miR-10b-5p was further corroborated by a global mouse knockout model. Specifically, congenital loss of miR-10b in mice resulted in body weight gain, increased susceptibility to diet-induced obesity, hyperglycaemia, impaired glucose tolerance and insulin resistance (107). Interestingly, miR-10b-5p mimic treatment reversed GI dysmotility, restored glucose tolerance and insulin sensitivity in transgenic mice where miR-10b-5p was knocked out, highlighting its superior long-term efficacy in treating both diabetes and GI dysmotility compared to commonly used antidiabetic and prokinetic medications (107,109).

In addition to its role in controlling metabolism, its relevance in the regulation of cell differentiation was previously described *in vitro*. In agreement with our study, the expression of miR-10b-5p was previously found to be significantly upregulated during adipogenesis in 3T3-L1 cells. At the molecular level, the activity of apolipoprotein L6 (Apol6), a lipid-binding protein that plays a key role in adipogenesis, was shown to be regulated by miR-10b-5p (103). On the contrary, conflicting results by Li et al. 2018 (110) showed that miR-10b increased during osteogenic differentiation but declined during adipogenic differentiation in human adipose-derived mesenchymal stem cells (hADSCs). Furthermore, overexpression of miR-10b was found to enhance osteogenesis and inhibit adipogenesis, with downregulation reversing these effects. ADSCs are commonly used to study adipogenesis, but their use is limited by the inherent heterogeneity of the population, which can vary based on the source and isolation method (111). This heterogeneity can lead to inconsistent results in adipogenesis studies, as it fails to account for differences between white and brown adipose tissue depots, potentially masking depot-specific variations in cell phenotype. Moreover, variations in culture techniques, media formulations, and adipogenic differentiation protocols can impact the consistency and reproducibility of results in adipogenesis studies. Further conflicting findings by Tan and colleagues (103) demonstrated that the downregulation of miR-10b-5p promoted the differentiation of 3T3-L1 cells and adipogenesis by upregulating the Apol6 expression. Their study achieved miR-10b knockdown or upregulation by treating cells every two days, raising the possibility that repeated exposure to transfection reagents influenced the differentiation process. This potential impact is reflected in their Oil Red O staining, where differentiation appears relatively low even at day 8. Importantly, Fabp4, a robust marker of adipogenesis (112), seems to be upregulated in response to both miR-10b mimic and inhibitor when compared to control. In contrast, our approach minimizes these variables, providing a clearer assessment of miR-10b role during adipogenesis.

There are still several limitations in this study. For instance, we only investigated the effect of miR-10b-5p in *in vitro* experiments and did not further verify the proposed mechanism *in vivo*. Secondly, the mature miR-10a-5p and miR-10b-5p are part of the miR-10 family, differing by just a single nucleotide in the middle of their sequences (113). These two miRNAs play a crucial role in diabetes regulation (109). miR-10a-5p was highly expressed in brown preadipocytes but downregulated upon differentiation, reaching levels similar to white mature adipocytes. Its expression in white adipocytes remained unchanged between preadipocytes and mature cells. This remains a limitation of the study, as it has not been specifically investigated. Nevertheless, Zogg et al. (107) showed that global knockout of miR-10b in a mouse model still resulted in metabolic syndrome symptoms, with only a slight increase in miR-10a-5p, suggesting that miR-10b-5p alone plays a critical role in the development of metabolic dysfunction.

## 8 Conclusion

This study shows that miR-10b inhibition plays a dynamic role in adipocyte biology, as its inhibitory effects manifest differently during the stem cell commitment state and the maturation phase of adipocytes. miR-10b-5p regulates early adipogenesis by guiding cells toward the mesoderm lineage through its regulatory role in the GATA6/BMP2 signalling axis, and later stages by downregulating Tub. Understanding the miR-10b-mediated regulatory mechanism during adipocyte commitment and differentiation may help to generate adipose tissue-engineering strategies for cellular therapies for lipodystrophy and obesity.

## 9 Abbreviations

AP: Alkaline Phosphatase
BA: brown adipocyte
DEGs: differentially expressed genes
ECAR: extracellular acidification rate
ES: embryonic stem
FACS: fluorescence-activated cell sorting
GO: Gene ontology
GO: Gene Ontology
GPCRs: G protein-coupled receptors
GSEA: Gene set enrichment analysis
HRMA: high resolution melt analysis
iBAT: interscapular brown adipose tissue
IPSC: Induced pluripotent stem cell
KO: knockout
LNA: locked nucleic acid
mESC: mouse embryonic stem cell
miRNA: microRNA
MSC: mesenchymal stem cell
NTC: non-targeted control
OCR: oxygen consumption rate
PCA: principal component analysis
scWAT: subcutaneous white adipose tissue
sgRNA: single guide RNA
SVF: stromal vascular fraction
TGs: triglycerides
TPM: transcripts per million
UTR: untranslated region
WA: white adipocyte
β3AR: beta 3 adrenergic receptor

## 10 Declarations

### 10.2 Author contribution

NK and MC conceived and designed the experiments and wrote the manuscript. NK performed the majority of experiments and data analysis. JH performed the immunofluorescence of BAT. AY performed the proliferation assay and run the Western blotting for scWAT time course. FD prepared the samples for miRNA PCR array assay. SC, AVP and CNG designed and performed the hESCs differentiation to BA. All authors contributed to editing the manuscript.

### 10.4 Consent for publication

Not applicable.

### 10.5 Ethics approval and consent to participate

Not applicable.

### 10.6 Availability of data and materials

The datasets used and/or analysed during the current study are available from the corresponding author on reasonable request. RNA sequencing datasets, including those from miR-10b^+/+^ and miR-10b^−/−^ mESCs during adipocyte differentiation at days 0, 12, and 27 (E-MTAB-15151); RNA sequencing analysis of miR-10b-5p knockdown during brown adipogenesis(E-MTAB-15152); and the identification of differentially expressed miRNAs between mouse scWAT and iBAT (E-MTAB-15230) are available at ArrayExpress.

## 10.7 Acknowledgements

The authors would like to thank Graham Hickman for his support with confocal microscopy.

