## Supplementary Materials for "miR-10b-5p regulates adipocyte lineage commitment and adipogenesis via targeting of Gata6 and Tubby"

### Supplementary Tables

#### *Supplementary Table 1. **Brown Fat Differentiation Protocol***

*Induction media was administered for 2 days. Following induction, cells were treated with maintenance medium for 6 to 8 days with medium being replenished every 2 days. Components were added in complete medium that included DMEM/F12, 10%FBS, 1% L-Glutamine and 1% Penicillin-Streptomycin antibiotics.*

| Compound | Induction Medium | Maintenance Medium |
|---|---|---|
| Insulin | 1 μg/ml | 1 μg/ml |
| 3,3′ ,5-Triiodo-L-thyronine sodium salt (T3) | 1 nM | 1 nM |
| IBMX | 0.5 mM | |
| Dexamethasone | 250 nM | |
| Indomethacin | 125 μM | |

#### *Supplementary Table 2. **White Fat Differentiation Protocol***

*Induction medium 1 was administered for 3 to4 days. Following induction medium 1, cells were treated with induction medium 2 for 2 to 3 days and then with maintenance medium for 2 to 3 days. Components were added in complete medium that included DMEM/F12, 10%FBS, 1% L-Glutamine and 1% Penicillin-Streptomycin antibiotics.*

| Compound | Induction Medium 1 | Induction Medium 2 | Maintenance Medium |
|---|---|---|---|
| Insulin | 5 mg/ml | 5 mg/ml | 5 mg/ml |
| IBMX | 0.5 mM | | |
| Dexamethasone | 250 nM | | |
| 3,3′,5-Triiodo-L-thyronine sodium salt (T3) | 0.1 nM | 0.1 nM | 0.1 nM |
| Cortisol | 100 nM | | |
| Transferrin | 1 mg/ml | 1 mg/ml | 1 mg/ml |
| Rosiglitazone | 5 mM | 1 mM | |

| | | | |
|---|---|---|---|
| Biotin | 16 mM | 16 mM | 16 mM |
| Panthothenic acid | 1.8 mM | 1.8 mM | 1.8 mM |
| Ascorbic acid | 100 mM | 100 mM | |

*Supplementary Table 3*. ***Primer sequences for human genes.***

| Gene name | Forward primer (5’ to 3’) | Reverse primer (5’ to 3’) |
|---|---|---|
| GATA6 | CACACCACAACTACCACCTTAT | TCCTGGTTTGAATTCCCTCTTT |
| FABP4 | AACTGGTGGTGGAATGCGT | AACTGGTGGTGGAATGCGT |
| LN19 | GCGGAAGGGTACAGCCAA | GCAGCCGGCGCAAAA |
| DLK1 | CACGGACTCTGTGGAGAACC | GCAGGCCCGAACATCTCTAT |
| UCP1 | GTGTGCCCAACTGTGCAATG | CCAGGATCCAAGTCGCAAGA |

*Supplementary Table 4*. ***Primer sequences for mouse genes.***

| Gene name | Forward primer (5’ to 3’) | Reverse primer (5’ to 3’) |
|---|---|---|
| BMP2 | GAACACAAGTCAGTGGGAGAG | CACCTGGGTTCTCCTCTAAATG |
| CIDEA | CACGCATTTCATGATCTTGGA | GTTGCTTGCAGACTGGGACAT |
| DAB2 | GACGCTTTCACTGGCTTAGA | CCTTCCTTGAGGGAACAAGAG |
| DLK1 | CGGGAAATTCTGCGAAATAG | TGTGCAGGAGCATTCGTACT |
| FABP4 | ACACCAGATTTCCTTAAACTG | CCATCTAGGGTTATGATGCTCTTCA |
| GATA4 | GGAAGCCCAAGAACCTGAATA | CTAGTGGCATTGCTGGAGTTA |
| GATA6 | GCCTTGTCTGCTAAGGAAGAT | GGATGAATGGGTTCTGGGATAA |
| KLF4 | GTGCCCCGACTAACCGTTG | GTCGTTGAACTCCTCGGTCT |
| LN19 | GAGCACATCCACAAGCTGAA | TTTCGTGCTTCCTTGGTCTT |
| NANOG | GCCTCCAGCAGATGCAAG | GGTTTTGAAACCAGGTCTTAACC |
| OCT4 | CGTGGAGACTTTGCAGCCTG | GCTTGGCAAACTGTTCTAGCTCCT |
| SOX7 | TCACCTCCCCCATCTACCAG | GGCCAAGGGCTAAAGAACCT |
| TUB | ACAATGGCGTCAACCCTCAG | CTGGGACGATCACACTCATCTTC |
| UCP1 | TACCCAAGCGTACCAAGCTG | ACCCGAGTCGCAGAAAAGAA |

*Supplementary Table 5*. ***Primer sequences for Luciferase and mutagenesis assays.***

| Gene name | Forward primer (5’ to 3’) | Reverse primer (5’ to 3’) |
|---|---|---|
| Gata6 3’UTR | GCTGGTGCTACCAAGAGGC | GGTTGGTCACGTGGTACAGG |
| Tub 3’UTR | GTTTCTAGAGGGCAGTAGG AC | GCCTGTCCTCACCAAGCTG |
| Gata6 3’UTR mutagenesis | AGAAAAATATCTTGTTTGCT ACCAGATTTACAAATTCCAA GTGACCTCAGATCAGCC | GGCTGATCTGAGGTCACTTGGAATT TGTAAATCTGGTAGCAAACAAGATAT TTTTCT |
| Tub 3’UTR mutagenesis | TAGAGATGACTGCTTAGCT AGGAAGCTCTGCTCTG | CAGAGCAGAGCTTCCTAGCTAAGCA GTCATCTCTA |

*Supplementary Table 6*. ***CRISPR/Cas9 sgRNA sequences without the NGG end and primers used for HRMA.***

| **Sequence name** | **Forward sequence (5’ to 3’)** | **Reverse sequence (5’ to 3’)** |
|---|---|---|
| miR-10b-5p sgRNA | CCTGTAGAACCGAATTTGTG | CACAAATTCGGTTCTACAGG |
| Non-Targeted Control sgRNA | GGGTCTTCGAGAAGACCT | AGGTCTTCTCGAAGACCC |
| HRMA primers | CCGAGGTTGTAACGTTGTC | CCATGTCGGAGATATATGAAG |

*Supplementary Table 7*. ***Oligonucleotide sequences of miRNA inhibitors and mimics.***

| **name** | **Catalogue number** | **Inhibitor sequence** |
|---|---|---|
| mmu-miR-10b-5p miRCURY LNA miRNA Inhibitor | YI04100556-DDA | ACAAATTCGGTTCTACAGGGT |
| miRCURY LNA miRNA Inhibitor Control | YI00199006 | TAACACGTCTATACGCCCA |
| **name** | **Catalogue number** | **Mature miRNA sequence** |
| hsa-miR-10b-5p miRCURY LNA miRNA Mimic | YM00472145 | UACCCUGUAGAACCGAAUUUGUG |
| Negative Control miRCURY LNA miRNA Mimic | YM00479902 | UCACCGGGUGUAAAUCAGCUUG |

Supplementary Table 8. **On/off target sites of sgRNA miR-10b generated by CCTop - CRISPR/Cas9 target online predictor.** (web tool accessed at 17:32pm, 07/12/23)

| Coordinates | MM | target_seq | PAM | position | gene name | gene id |
|---|---|---|---|---|---|---|
| chr2:74726077-74726099 | 0 | CCTGTAGA[ACCGAATTTGTG] | TGG | Exonic | Mir10b | ENSMUSG00000065500 |
| chr2:105460157-105460179 | 4 | TATGTGGA[AACGAATTTGTG] | AGG | Intergenic | 4930527A07Rik | ENSMUSG00000086764 |
| chr15:59583720-59583742 | 4 | ATTGGAGA[ACTGAATTTGTG] | TGG | Intronic | Nsmce2 | ENSMUSG00000059586 |
| chr5:109894569-109894591 | 4 | TCTTTAAA[ACGGAATTTGTG] | GGG | Intergenic | Gm26779 | ENSMUSG00000097140 |
| chr9:41431921-41431943 | 4 | GCTGAACA[ACTGAATTTGTG] | GGG | Exonic | 2610203C20Rik | ENSMUSG00000074415 |

| | | | | | | |
|---|---|---|---|---|---|---|
| chr1:58718061-58718083 | 4 | GCTGCAGT[ACTGAATTTGTG] | GGG | Intronic | Cflar | ENSMUSG00000026031 |
| chr8:106690664-106690686 | 4 | TCTGGAGC[ACAGAATTTGTG] | TGG | Intronic | Tango6 | ENSMUSG00000041949 |
| chr4:40339598-40339620 | 4 | TCTATAGA[GCTGAATTTGTG] | AGG | Intergenic | 4930509K18Rik | ENSMUSG00000087137 |
| chr15:72912507-72912529 | 4 | AGTGTAGA[AGCTAATTTGTG] | AGG | Intronic | Gm3150 | ENSMUSG00000096173 |
| chr16:84708197-84708219 | 3 | CTTGTAGA[ACTTAATTTGTG] | TGG | Intronic | Mir155hg | ENSMUSG00000097418 |
| chr18:40483559-40483581 | 4 | ACTGTAGT[CCTGAATTTGTG] | GGG | Intronic | Kctd16 | ENSMUSG00000051401 |
| chr5:16220572-16220594 | 4 | CCTTCAGA[AATGAATTTGTG] | TGG | Intronic | Cacna2d1 | ENSMUSG00000040118 |
| chr18:29833061-29833083 | 4 | TCTTTAGA[AACCAATTTGTG] | CGG | Intergenic | NA | NA |
| chr19:42216214-42216236 | 4 | CCTTTAGT[ATCTAATTTGTG] | TGG | Intergenic | Sfrp5 | ENSMUSG00000018822 |
| chr17:3969757-3969779 | 4 | CATTTACA[ACCGAAATTGTG] | AGG | Intergenic | NA | NA |
| chr6:36427248-36427270 | 4 | ACAGTAGA[ACAGATTTTGTG] | TGG | Intronic | Mir490 | ENSMUSG00000070075 |
| chr5:124266586-124266608 | 4 | CTTTTAGA[ACCTAGTTTGTG] | TGG | Intronic | Mphosph9 | ENSMUSG00000038126 |
| chr11:107860036-107860058 | 4 | TCTGCAGA[AGCGAATGTGTG] | TGG | Intergenic | Gm27595 | ENSMUSG00000098991 |
| chr12:112107864-112107886 | 3 | CCTGTTGA[ACCTAATTGGTG] | TGG | Intronic | Aspg | ENSMUSG00000037686 |
| chr15:3327419-3327441 | 4 | CCTGAACA[ACAGAATTTCTG] | GGG | Intronic | Ghr | ENSMUSG00000055737 |

*Supplementary Table 9*. ***Study design of the transcriptomic analysis performed on mESCs differentiation to mature adipocytes.***

| Day 0 | | Day 12 | | Day 27 | |
|---|---|---|---|---|---|
| **Clone** | **Number of samples** | **Clone** | **Number of samples** | **Clone** | **Number of samples** |
| NCT Clone 1 | 3 | NCT Clone 1 | 1 | NCT Clone 1 | 3 |
| NCT Clone 2 | 1 | NCT Clone 2 | 2 | NCT Clone 2 | |
| NCT Clone 3 | 1 | NCT Clone 3 | 2 | NCT Clone 3 | 1 |
| 1F10 | 3 | 1F10 | 3 | 1F10 | 2 |
| 3G6 | 1 | 3G6 | 1 | 3G6 | |
| 3F2 | 1 | 3F2 | | 3F2 | 1 |

### Supplementary figures

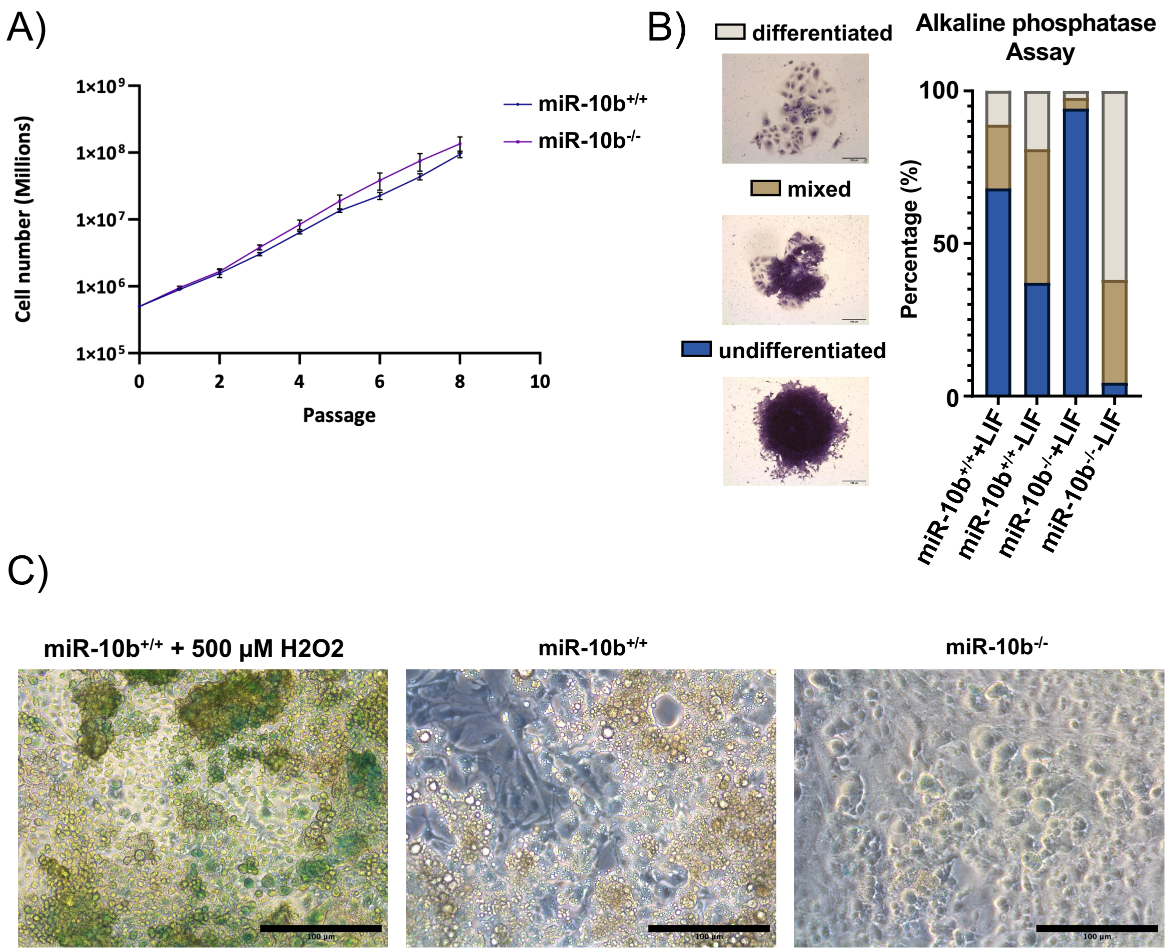


*Figure S1*. ***Assessment of mESC dynamics with/without miR-10b: proliferation, senescence, and alkaline phosphatase activity analysed.*** *A) Cell proliferation*

*assay. Data are representative of three independent experiments and values are expressed in mean ± SEM. B) Alkaline Phosphatase (AP) activity was used to assess the self-renewal capacity of mESCs. Scale bars: 100 µm. C) Cell senescence was analysed in cells with or without depleted miR-10b. miR-10b+/+ and miR-10b-/- stem cells were differentiated to mature adipocytes. At day 27, cells were stained for β-galactosidase and observed at ×20 magnification. For positive control, miR-10b-/- were treated with 500 µM H2O2 for 1 hour. Scale bar represents 100 µM. β-galactosidase-positive cells appear light blue.*

A)

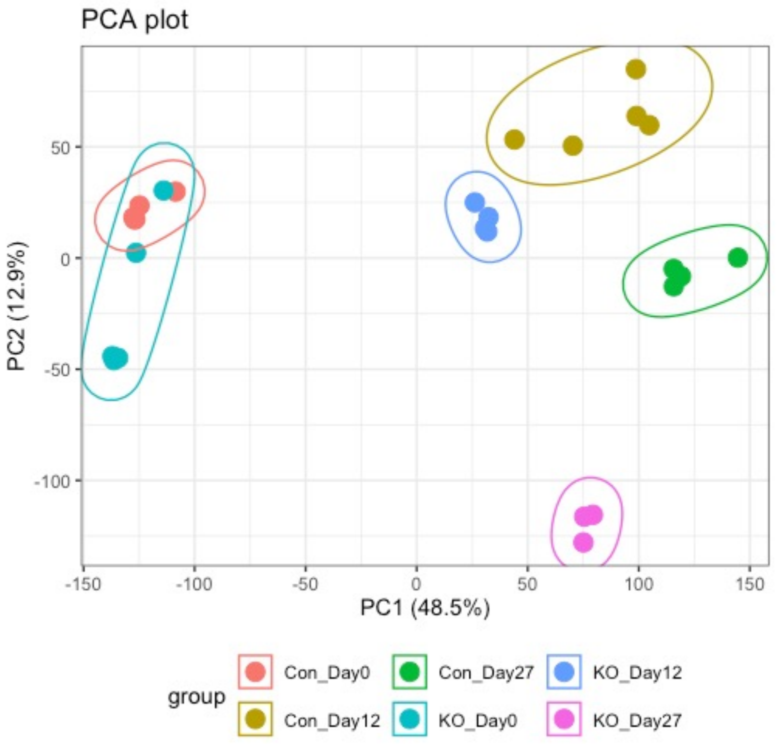


B)

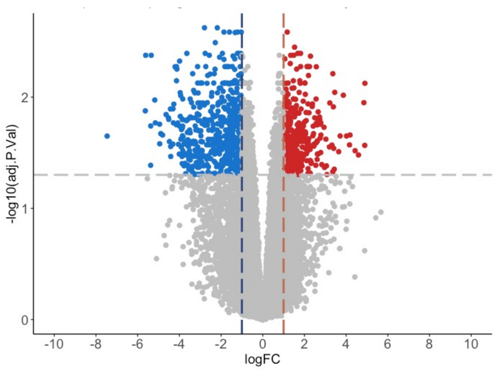


*Figure S2*. ***Transcriptomic analysis was conducted on mESCs differentiating to mature adipocytes at day 0, 12, and 27.*** *A) PCA plot for stem cells time course. B) Volcano plot illustrating the expression profile of genes in mESCs treated with gRNA targeting miR-10b or NTC vector at Day 0 (log2(FC) > 1 and p adjusted value > 0.05).*

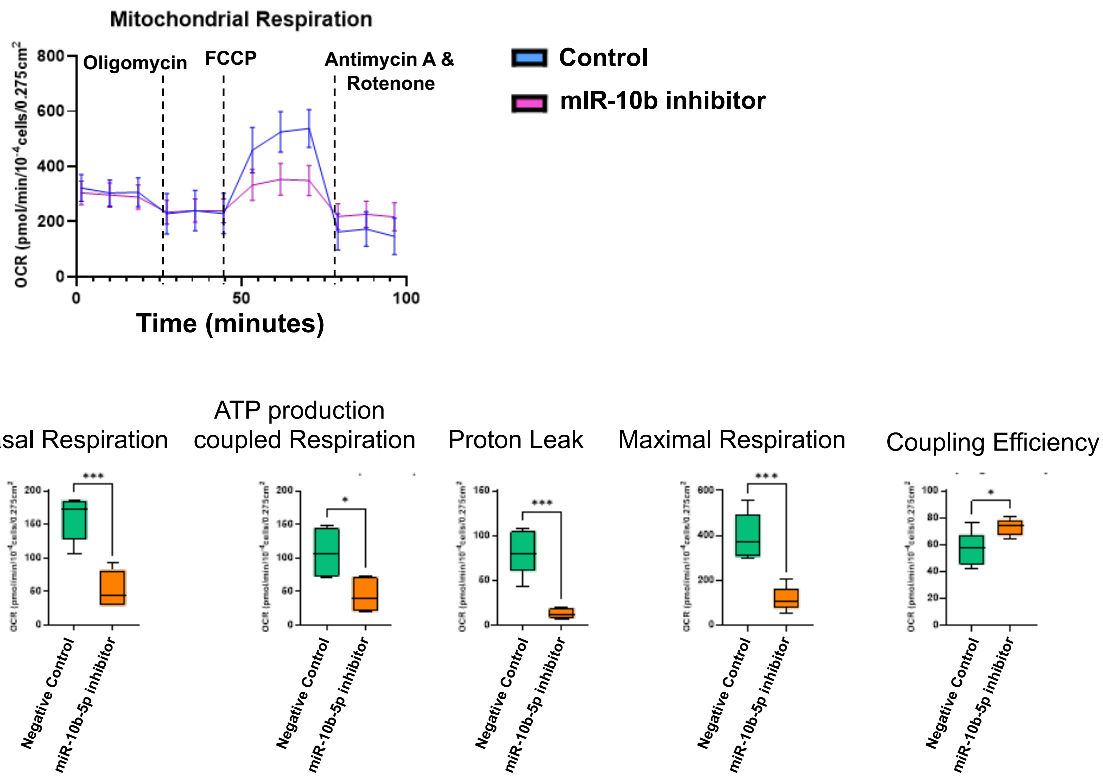


*Figure S3*. ***Functional effects of miR-10b Inhibition on Bioenergetics.*** *Brown preadipocytes were differentiated to mature BA for 7 days and then transfected with miR-10b-5p inhibitor or negative control inhibitor. Extracellular acidification rate (ECAR) and oxygen consumption rate (OCR) were assessed 48 hours post-transfection (n ≥ 5). The blue line represents the negative control, while the pink line corresponds to the miR-10b-5p inhibitor. Box and whisker plots illustrate changes in mitochondrial respiration components in differentiated BA, including ATP production-coupled respiration, proton leak, and maximal respiratory capacity.*

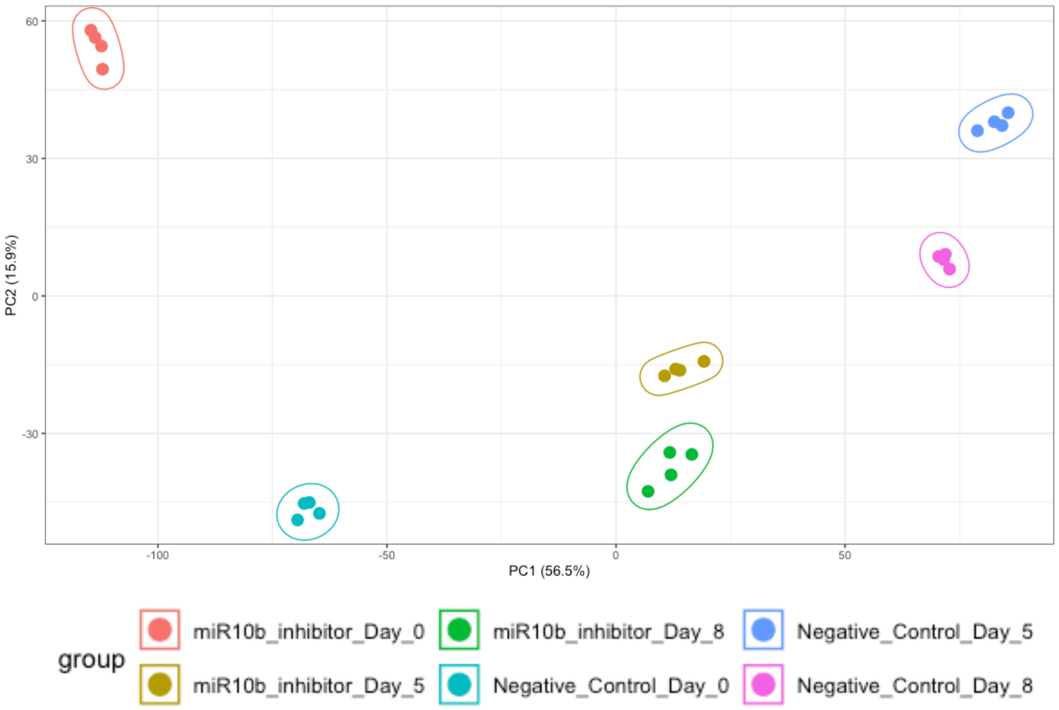


*Figure S4*. ***PCA plot for BAT time course.***

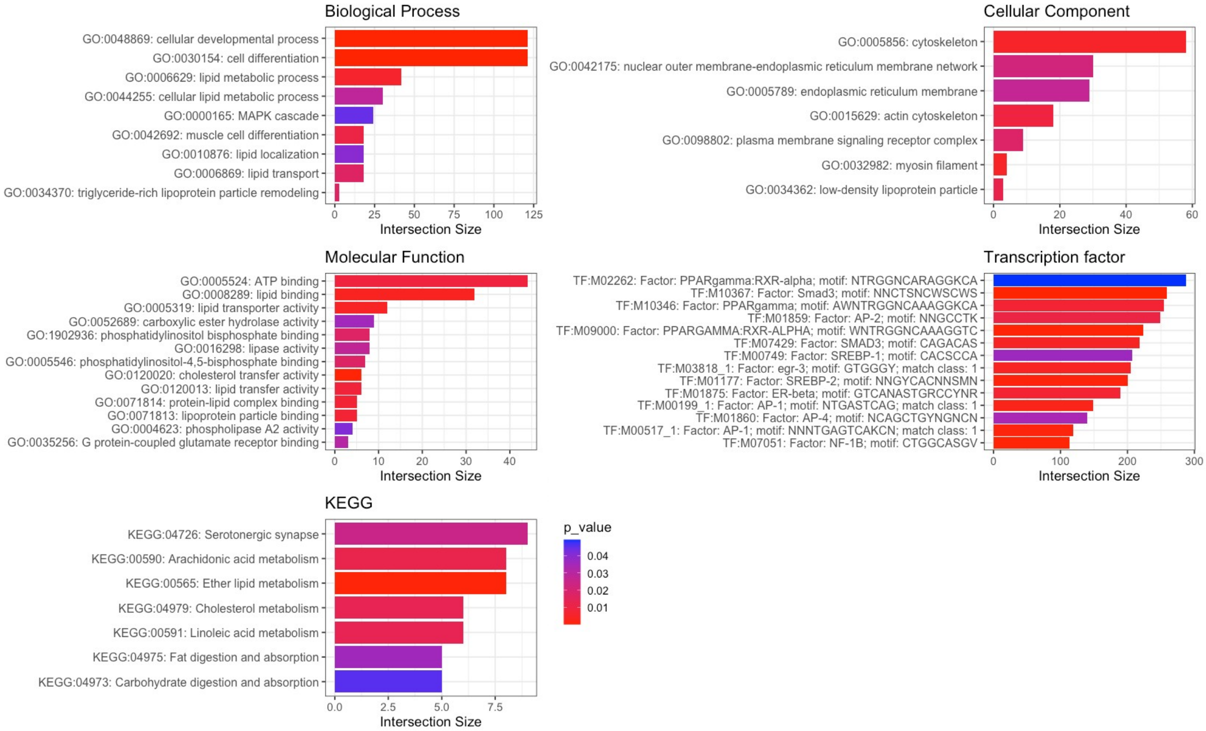


*Figure S5*. ***Gene ontology (GO) enrichment analysis of related differentially expressed genes (DEGs) in purple module.***

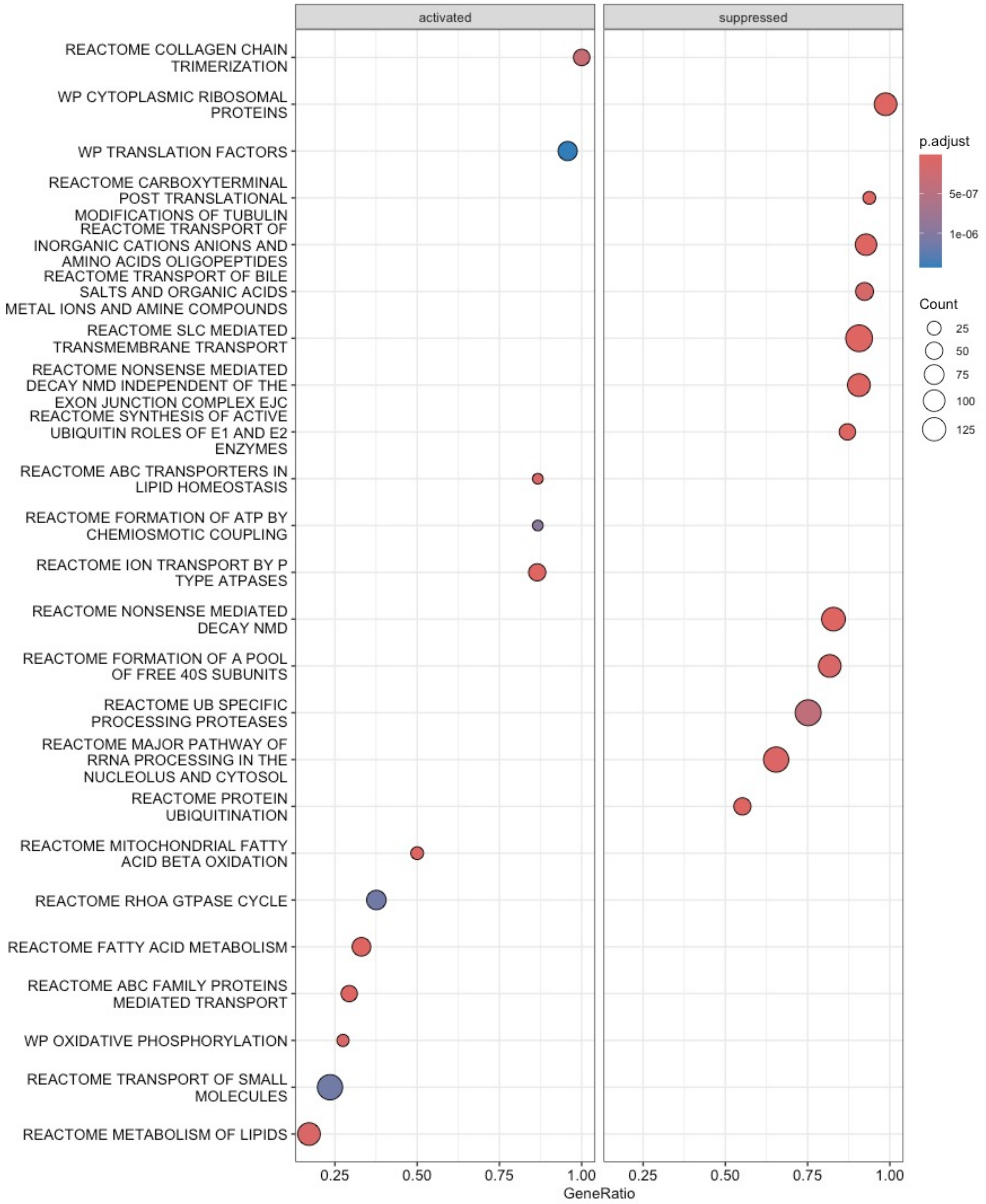


*Figure S6*. ***Dot plot of top 12 enriched pathways in DEGs from miR-10b-inhibited brown adipocytes, Day 0.*** *The count represents the number of inputted DEGs as a percentage of the total number of genes. The Benjamini p value for each molecular mechanism is shown.*

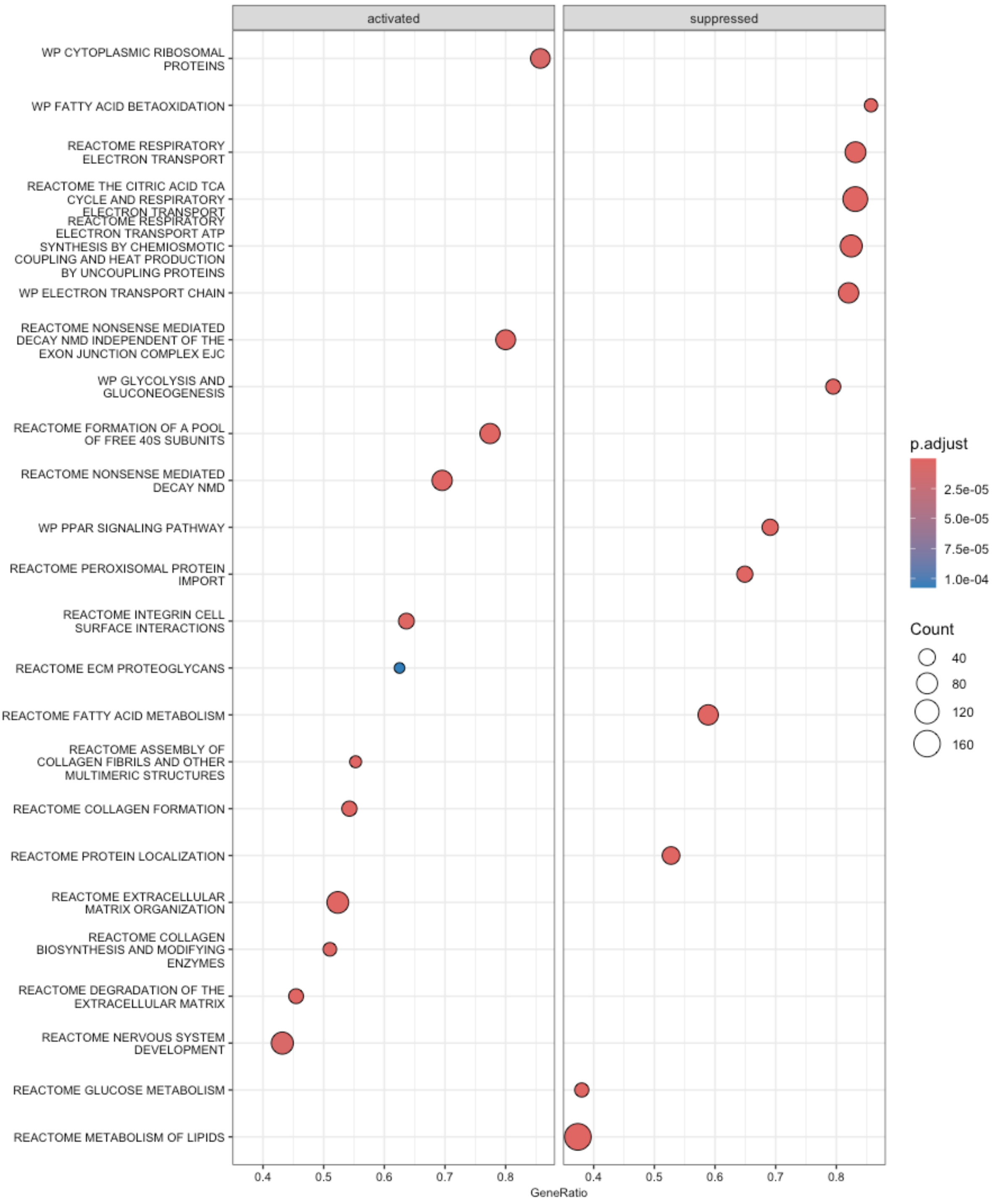


*Figure S7*. ***Dot plot showing top 12 enriched pathways in DEGs from miR-10b-inhibited brown adipocytes, Day 5.*** *Each dot represents the percentage of DEGs within the total number of genes. Additionally, the Benjamini-adjusted p-value for each molecular mechanism is presented.*

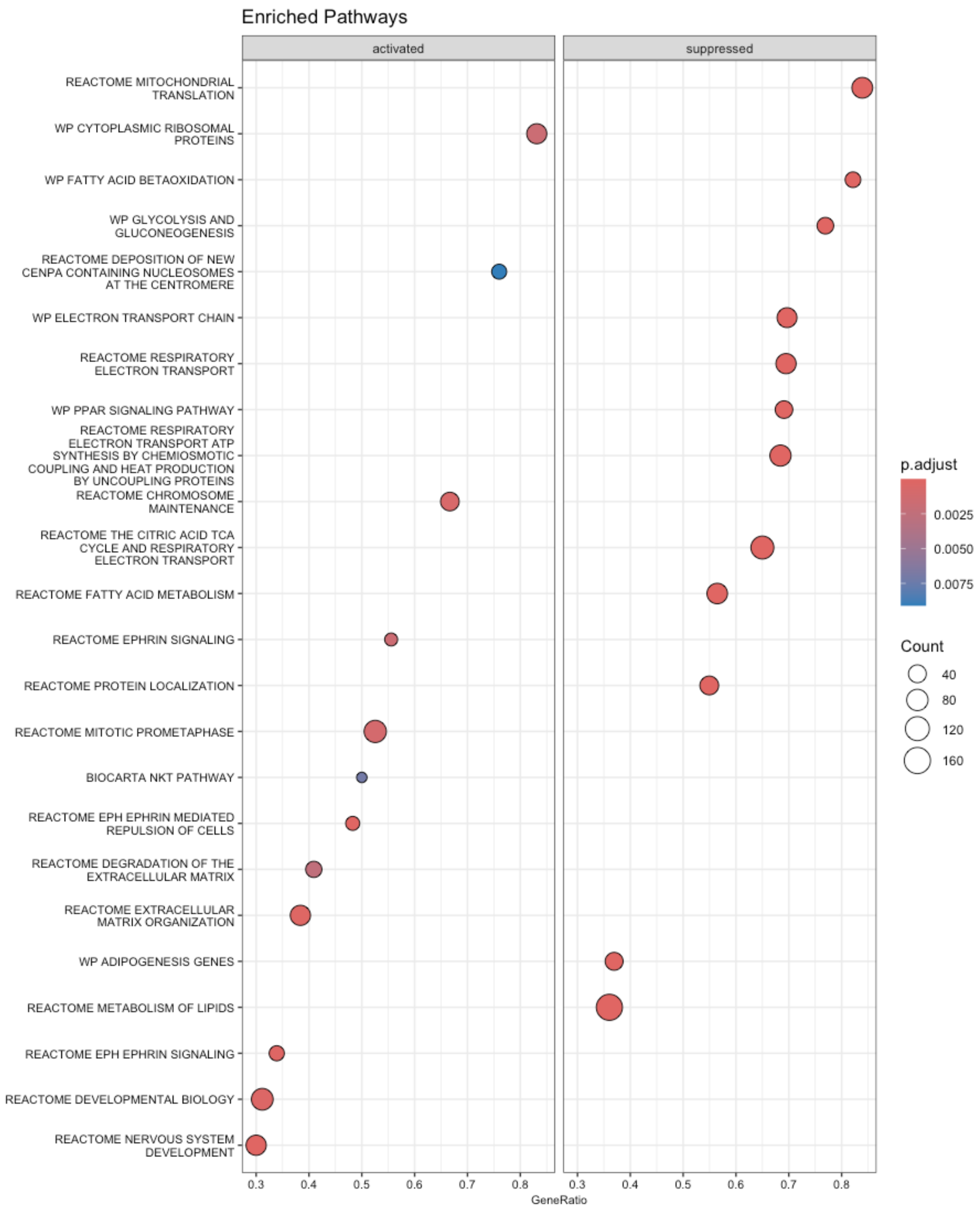


*Figure S8*. ***Dot plot showing top 12 enriched pathways in DEGs from miR-10b-inhibited brown adipocytes, Day 8.*** *The dots represent the percentage of DEGs within each pathway relative to the total number of genes in that category. Additionally, the Benjamini p-value for each pathway is displayed to indicate its statistical significance.*

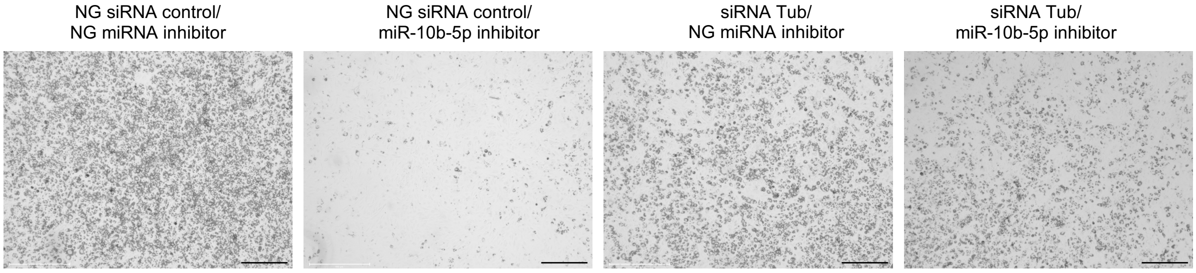


*Figure S9*. ***Bright-Field Imaging of Lipid Droplet Formation on Day 6 of Adipocyte Differentiation.*** *Representative 4× magnification images show lipid droplet accumulation as an indicator of differentiation on Day 6. Scale bar: 500 µm (black).*

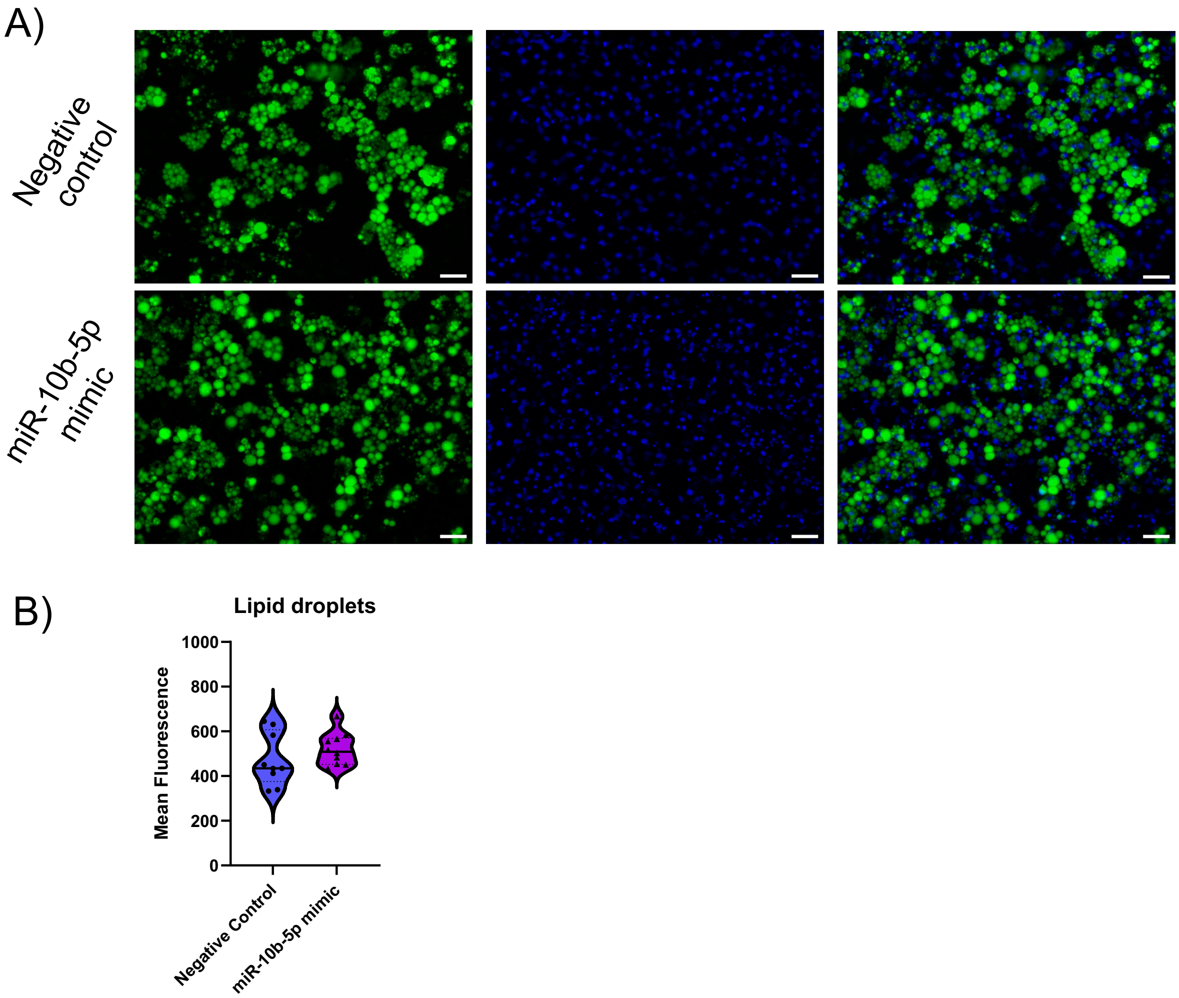


*Figure S10*. ***The effects of elevated miR-10b-5p levels on white adipogenesis.*** *A) White preadipocytes were treated with either negative control or miR-10b-5p mimic for 48 hours and differentiated for 8 days, followed by staining of their lipids using a GFP lipid stain and their nucleus with DAPI. Scale bar: 50 B) Lipid droplet quantification.*

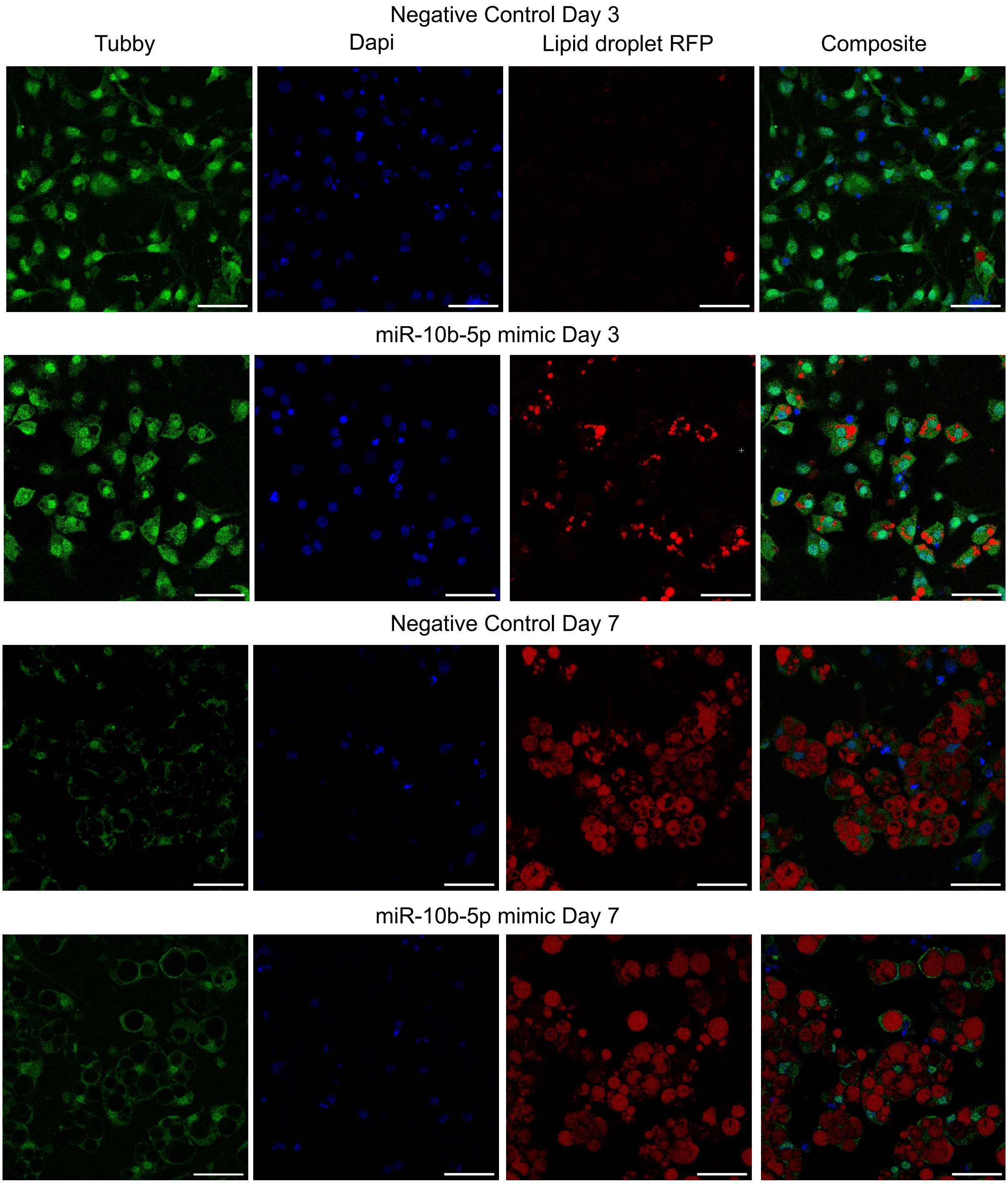


*Figure S11*. ***Tubby immunostaining during WAT differentiation. Scale bar: 50 µm.***

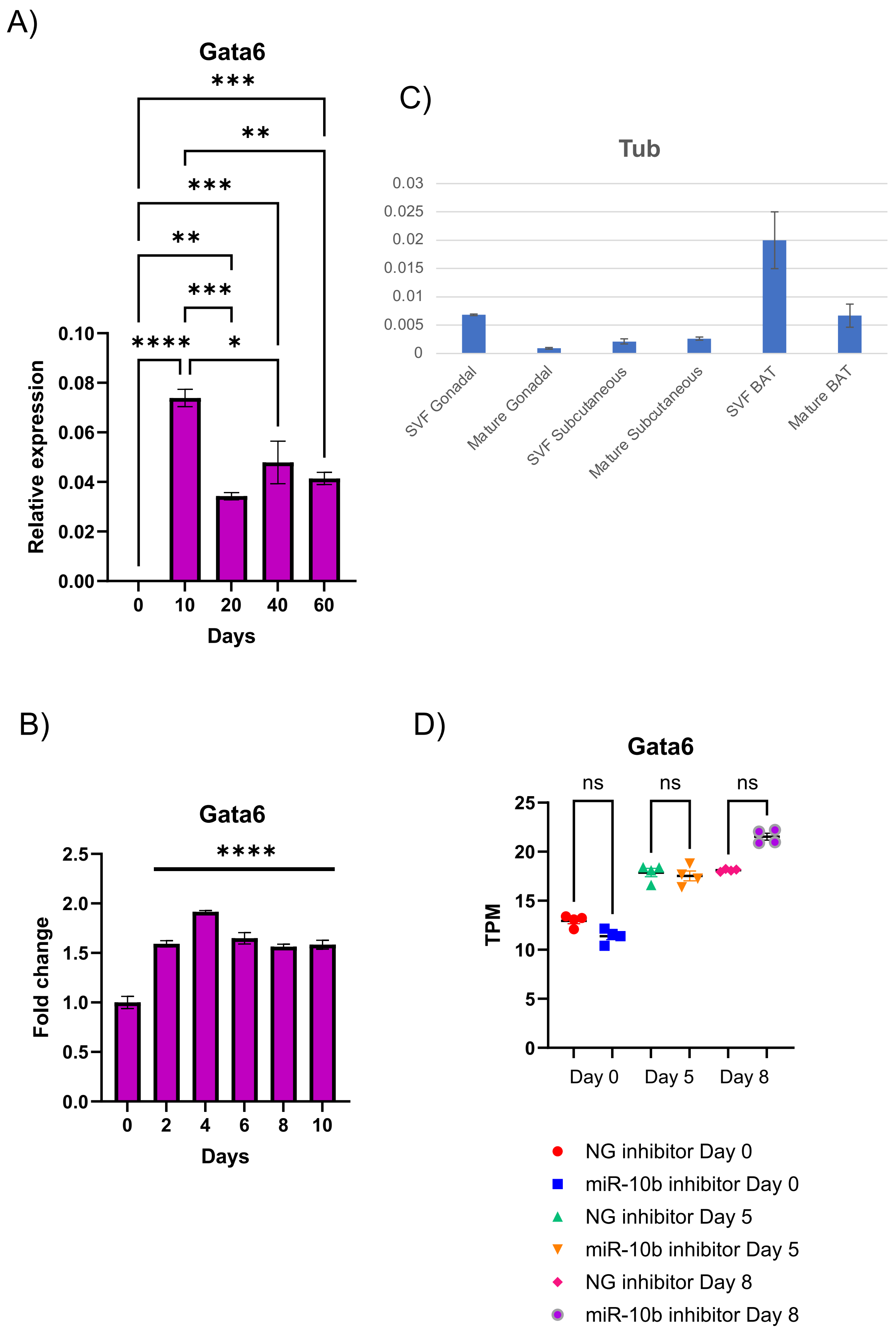
A)
Gata6
Relative expression
0.00
0.02
0.04
0.06
0.08
0.10
0
10
20
40
60
Days
****
*
***
**
***
**
***
C)
Tub
0
0.005
0.01
0.015
0.02
0.025
0.03
SVF Gonadal
Mature Gonadal
SVF Subcutaneous
Mature Subcutaneous
SVF BAT
Mature BAT
B)
Gata6
Fold change
0.0
0.5
1.0
1.5
2.0
2.5
0
2
4
6
8
10
Days
****
D)
Gata6
TPM
0
5
10
15
20
25
ns
ns
ns
Day 0
Day 5
Day 8
NG inhibitor Day 0
miR-10b inhibitor Day 0
NG inhibitor Day 5
miR-10b inhibitor Day 5
NG inhibitor Day 8
miR-10b inhibitor Day 8

*Figure S12*. ***Gata6 expression during human and mouse brown adipocyte differentiation and across various adipose tissue depots.*** *A) Gata6 expression during human ESC differentiation to BA. Data ($n \geq 3$) are shown as means + SEM (two-way ANOVA test, *$p < 0.05$, **$p < 0.005$, ***$p < 0.0005$, ****$p < 0.0001$). B) Gata6 expression during mouse BA differentiation. Data ($n \geq 3$) are shown as means + SEM (two-way ANOVA test, ****$p < 0.0001$). C) Tub expression across different adipose tissue depots. D) TPM levels of Gata6 expression during mouse BA differentiation, as measured by RNA sequencing. Data ($n \geq 3$) are shown as means + SEM (Kruskal-Wallis test, ns: not significant).*
